# Endothelial Sphingosine-1-Phosphate Receptor (S1PR) 1, a Functional S1PR in the Human Cerebrovascular Endothelium, Limits Blood Brain Barrier Permeability and Neuronal Injury following Subarachnoid Hemorrhage in Mice

**DOI:** 10.1101/2025.01.19.633813

**Authors:** Akira Ito, Hiroki Uchida, Sabyasachi Dash, Daiki Aburakawa, Yuto Shingai, Cathy Liu, Paulo Avila-Gomez, Keita Tominaga, Gab Seok Kim, Giuseppe Faraco, Richard L. Proia, Kuniyasu Niizuma, Teiji Tominaga, Josef Anrather, Costantino Iadecola, Michael J. Kluk, Teresa Sanchez

## Abstract

Hypoxia-induced blood-brain barrier (BBB) permeability has been identified as a key contributor to the progression of ischemic-hypoxic brain injury and neuronal dysfunction in stroke and other cerebrovascular diseases. Emerging clinical evidence highlights that the vasoprotective signaling properties of high-density lipoprotein (HDL), mediated through its bioactive lipid component sphingosine-1-phosphate (S1P), may be impaired in cardiovascular and inflammatory conditions. Nonetheless, the precise contributions and mechanistic roles of S1P signaling within the cerebral microvasculature remain insufficiently characterized.

In this study, we aimed to elucidate the role of S1P signaling via its endothelial receptor S1PR1 in the pathophysiology of early brain injury following subarachnoid hemorrhage (SAH), a particularly severe form of stroke. Additionally, we sought to evaluate the relevance of the endothelial S1PR1 pathway in the human cerebrovascular endothelium, its functional role in hypoxia-induced cerebral endothelial barrier dysfunction, and its underlying molecular mechanisms. To address these objectives, we utilized endothelial-specific S1PR1 knockout mice subjected to the endovascular rupture model of aneurysmal SAH, performed mechanistic studies in primary human cerebral microvascular endothelial cells, and characterized S1PR1 expression in human brain tissue using validated protocols.

Our findings reveal robust expression of S1PR1 in the cerebrovascular endothelium of both mice and humans. Functional analyses demonstrated that S1PR1 is critical for maintaining BBB integrity and mitigating neuronal injury in the context of SAH. Mechanistic *in vitro* studies indicated that S1PR1 exerts a vasoprotective effect by limiting hypoxia-induced BBB dysfunction in human primary brain microvascular endothelial cells through inhibition of Rho-associated kinase (ROCK)-mediated phosphorylation of myosin light chain (MLC), suppression of stress fiber formation and caveolin-1-dependent endosomal trafficking.

These results highlight the pivotal role of endothelial S1PR1 signaling in preserving cerebral vascular integrity and provide a strong scientific foundation for developing novel therapeutic approaches targeting the S1P pathway in the endothelium to enhance neurovascular protection.

## INTRODUCTION

Cerebrovascular disease represents a leading cause of death and disability world-wide^1^ ^2^ ^3^ and is a significant contributor to stroke and dementia^4^. Aneurysmal SAH, the most devastating type of stroke, results from the rupture of an intracranial aneurysm. Compared to other types of stroke, SAH occurs earlier in life (typically between 40-60 years of age)^5^ and is associated with higher mortality (∼50%) and greater long-term disability^6^. Cerebral aneurysm rupture leads to accumulation of blood in the subarachnoid space, transient cerebral ischemia followed by hypoperfusion and subsequent hypoxic brain injury^7, 8^, inflammation, cerebral edema, and blood-brain barrier (BBB) dysfunction ^9–13^. While recent advances in surgical interventions for aneurysm repair have reduced mortality, there are currently no effective pharmacological strategies to mitigate the extensive microvascular damage and early brain injury ^14^ caused by SAH. Consequently, survivors experience a high degree of disability, motor and cognitive impairments (e.g., memory, language and executive function) ^15^ which impose substantial personal and societal burdens. Mounting preclinical and clinical evidence implicates BBB dysfunction in the pathophysiology of early brain injury following SAH ^11, 16–21^ ^12^ ^22, 23^. The cerebrovascular endothelium, strategically located at the interface between the bloodstream and brain parenchyma and as a pivotal regulator of BBB integrity and neurovascular homeostasis, constitutes a promising therapeutic opportunity. However, the endothelial signaling pathways and molecular mechanisms that govern hypoxia-ischemia-induced BBB dysfunction remain poorly understood, creating a significant barrier to the development of targeted endothelial therapies for mitigating neurovascular injury in SAH and related hypoxic-ischemic cerebrovascular conditions.

The sphingosine-1-phosphate (S1P) pathway is a cornerstone of vascular^24^ and immune^25^ homeostasis in both humans and mice, and is gaining recognition as an attractive target for drug development. S1P, a bioactive sphingolipid primarily produced by endothelial cells and erythrocytes, is an integral component of high-density lipoproteins (HDL) ^26, 27^. Within the endothelium, S1P exerts its effects through G-protein-coupled receptors (S1PRs), particularly S1PR1, which activates the vasoprotective phosphatidylinositol-3-kinase (PI3K)-Akt-eNOS signaling cascade^28^ ^29, 30^ ^31^. This pathway opposes the proinflammatory Rho-Rho kinase (ROCK) signaling axis^32–39^. S1PR1 enhances endothelial barrier function ^28, 40^ ^29, 30^ ^31, 41^ *by modulating* cytoskeletal dynamics and promoting adherens junction assembly ^29, 40, 42^, mitigates endothelial inflammation ^28, 42^ and mediates some of the cardiovascular protective effects of HDL ^31, 42^. Recent human and mouse studies indicate that vasoprotective HDL signaling via S1PR1 may be limited in cardiovascular and inflammatory diseases. While the functions of S1PR1 in the cardiovascular system are well characterized, its role in the cerebral microvasculature remains insufficiently understood.

Beyond its role in the vascular system, S1P-S1PR1 signaling is also essential in the immune system, regulating lymphocyte trafficking and function^25^. S1PR1 functional antagonists, such as Fingolimod (FTY720) ^43, 44^ and Siponimod (BAF312) ^45^, are already clinically approved as immunosuppressants for the treatment of multiple sclerosis. Interest in these agents is expanding to include cerebrovascular diseases and stroke owing to their protective effects in experimental models of ischemic stroke ^46, 47, 48^ ^49^ and subarachnoid hemorrhage ^50, 51^, largely attributed to immunosuppression ^48^ and S1PR1 blockade (reviewed in ^30^). Notably, pilot clinical trials suggest that FTY720 improves stroke outcomes^52–54^, underscoring the importance of immune modulation via S1PR1 inhibition. However, these findings leave the specific role of S1PR1 in the human cerebrovascular endothelium in BBB regulation during hypoxic-ischemic injury and its contributions to brain injury unresolved, representing a critical knowledge gap in the development of endothelial-targeted therapies. Further investigation is required to inform the design of innovative therapeutic strategies targeting this pathway in the cerebrovascular endothelium to prevent hypoxic-ischemic brain injury while minimizing the risks of immune suppression, which can have fatal consequences in critically ill and elderly patients ^30^.

In this study, we sought to address these critical unresolved questions by using a combination of genetic *in vivo*, *in vitro*, and translational approaches. We investigated the role of endothelial-specific S1PR1 in the pathophysiology of early brain injury following SAH. Additionally, we explored the potential relevance of the endothelial S1PR1 pathway in humans by examining the expression of S1PR1 in the cerebrovascular endothelium of the human brain, its functional role in hypoxia-induced barrier permeability *in vitro*, and the underlying molecular mechanisms. Our findings reveal that endothelial S1PR1 exerts a protective role in limiting BBB dysfunction and brain injury, identifying a previously unrecognized function of the endothelial S1PR1 pathway in the pathophysiology of early brain injury in SAH. We also demonstrate that S1PR1 is prevalent in the human cerebrovascular endothelium and functions as an endogenous protective signaling pathway *in vitro* mitigating hypoxia-induced endothelial barrier dysfunction in human cerebral microvascular endothelial cells. Mechanistically, we found that S1PR1 limits Rho-ROCK mediated cellular processes that lead to barrier leakage, including cytoskeletal remodeling (e.g., phosphorylation of myosin light chain and formation of stress fibers) and caveolin-1 trafficking to endosomal vesicles. These findings highlight the relevance of the endothelial S1PR1 pathway in protecting against BBB dysfunction and brain injury in SAH and other hypoxic-ischemic conditions. They provide a compelling rationale for developing innovative therapeutic strategies to specifically target S1P signaling in the endothelium for the treatment of cerebrovascular diseases while minimizing the risks associated with systemic immunosuppression of current S1PR1-modulating therapies ^30^.

## METHODS

### Mice

All animal experiments were approved by the Weill Cornell Institutional Animal Care and Use Committee. Endothelial cell specific *S1pr1* knockout mice (*S1pr1^flox/flox^xCdh5–Cre^ERT2^*; referred to as *S1pr1^iECKO^*) were generated as we have described ^55^. *S1pr1^flox/flox^*mice ^56^ were crossed to *Cdh5–Cre^ERT2^*mice ^57^ to generate *S1pr1^flox/flox^ Cdh5-Cre^ERT2^* mice. Mice were treated with tamoxifen (Sigma-Aldrich) by oral gavage (75 mg kg^-1^) for 3 days at the age of 8 weeks and used for the experiments 3-4 weeks after tamoxifen treatment. *S1pr1^flox/flox^* littermates treated with tamoxifen were used as control mice. S1pr1-eGFP knock in mice ^58^ weighing 26-30 g were used for the expression studies. All experiments were performed in male mice.

### Isolation of cortical microvessels

The brain microvessels were isolated as we have previously described ^59^. All procedures were performed in a cold room. The brains were collected and rinsed in MCDB131 medium (Thermo Fisher Scientific) with 0.5% fatty acid-free BSA (Millipore Sigma). The leptomeninges, cerebellum, brainstem and white matter were removed on ice. Ipsilateral cortices were homogenized in 8 mL of MCDB131 medium with 0.5% fatty acid-free BSA using a 7-mL loose-fit Dounce tissue grinder (Sigma-Aldrich) with 10 strokes. The homogenates were centrifuged at 2,000 g for 5 min at 4 °C. The pellet was suspended in 15% dextran (molecular weight ∼70,000 Da, Sigma-Aldrich) in PBS and centrifuged at 10,000 g for 15 min at 4 °C. The pellet was resuspended in MCDB131 with 0.5% fatty acid-free BSA and centrifuged at 5,000 g for 10 min at 4 °C. The pellet contained the brain microvessels.

### Endovascular perforation SAH surgery

SAH surgery was performed on C57BL6/J, *S1pr1^iECKO^* mice and their littermate *S1pr1^flox/flox^*controls as we have previously described ^60^. In brief, surgery was performed using a dissecting surgical microscope. Temperature was maintained at 36.5–37.5 °C by using a thermostatic blanket (Harvard Apparatus, CMA 450 Animal Temperature Controller) throughout the procedure. Mice were anesthetized with isoflurane inhalation delivered by facemask with O_2_. A 15 mm midline vertical incision was made on the skin in the head. The head was fixed in stereotactic frame and cerebral blood flow was measured by laser speckle imager (PeriCam PSI system, Perimed, Sweden). During surgery, mice were in supine position. A 10 mm midline vertical incision was made on the skin in the neck. The common carotid, external carotid and internal arteries were dissected from the adjacent tissue. The superior thyroid artery and the occipital artery were cauterized and cut. The external carotid artery was sutured with a dead knot and cauterized above the suture. A second suture loop was also placed in the external carotid artery just before the bifurcation of the common carotid artery. A slit-knot was placed around the common carotid artery. A small clip was applied to the internal carotid artery and the slip-knot around the common carotid artery was tightened temporally. A small incision was made in the external carotid artery stump. In young mice, a 5-0 monofilament with a modified tip (0.3 mm x 0.3 mm) was inserted into the incision and the knot around the external carotid artery was tightened to prevent bleeding. In middle-aged mice (∼11-12-month-old) an unmodified 5-0 monofilament was used to keep mortality <50%. Then, the monofilament was advanced to the common carotid artery, the small clip on the internal carotid artery was removed and the monofilament was guided through the external carotid artery to the internal carotid artery. The knot around the common carotid artery was opened again and the monofilament was introduced to the bifurcation of the internal carotid artery. The monofilament was gently pushed ∼1 mm further and then withdrawn to the external carotid artery. The knot around the external carotid artery was loosen and the monofilament was slowly removed. The external carotid artery was quickly ligated to prevent bleeding. The mouse was turned in prone position and induction of subarachnoid hemorrhage was confirmed by reduction of cerebral blood flow by laser speckle contrast imager. After the surgery, all animals were maintained in a small animal heated recovery chamber. The surgeon and the investigator conducting the analysis were blinded to the genotype of the mice. Animals which did not exhibit a reduction in CBF upon endovascular rupture were excluded from the study.

#### RNA isolation, reverse transcription and quantitative PCR analysis (RT-qPCR)

Total RNA from mouse brain, cells and microvessels was prepared using RNeasy Mini Kit (Qiagen, Valencia, CA) as instructed by the manufacturer. To generate cDNA, 100 ng of RNA was reverse transcribed using random primers and SuperScript II RT-polymerase (Invitrogen, Carlsbad, CA). Primers were designed using the Primer Express oligo design program software (Applied Biosystems, Foster City, CA). Real-time quantitative PCR was performed using the SYBR Green I assay on the ABI 7500 Sequence Detection System (Applied Biosystems). PCR reactions for each cDNA sample were performed in duplicate and copy numbers were calculated using standard curves generated from a master template as we previously described ^61^. The sequence of the primers used for qPCR are shown in Table 2.

**Table 1.** Physiological variables in *S1pr1^flox/flox^* and *S1pr1^iECKO^* mice.

|  |  |  |  |
| --- | --- | --- | --- |
| Time points | Parameters | *S1pr1<sup>flox/flox</sup>* | *S1pr1<sup>iECKO</sup>* |
| 5 min before SAH | Arterial O<sub>2</sub> saturation (SpO<sub>2</sub>, %) | 98.6 ± 0.2 | 98.4 ± 0.1 |
|  | Heart rate (b.p.m.) | 442.0 ± 23.2 | 459.2 ± 10.7 |
|  | Pulse distention (μm) | 13.8 ± 0.7 | 12.8 ± 0.9 |
|  | Respiratory rate (br.p.m.) | 162.3 ± 4.9 | 155.0 ± 6.7 |
| 5 min after SAH | Arterial O<sub>2</sub> saturation (SpO<sub>2</sub>, %) | 98.6 ± 0.1 | 98.6 ± 0.2 |
|  | Heart rate (b.p.m.) | 539.8 ± 9.9 | 549.7 ± 9.9 |
|  | Pulse distention (μm) | 13.2 ± 1.1 | 12.5 ± 0.9 |
|  | Respiratory rate (br.p.m.) | 133.3 ± 4.4 | 126.8 ± 6.5 |
| bpm, beats per minute; br.p.m., breaths per minute. Pulse distention is a surrogate for pulse pressure (pulse pressure 1/4 systolic pressure diastolic pressure). No significant differences were observed in the physiological parameters (arterial O<sub>2</sub> saturation, heart rate, pulse distention and respiratory rate) between *S1pr1<sup>flox/flox</sup>* and *S1pr1<sup>iECKO</sup>* mice, before and after SAH |  |  |  |

**Table 2.**
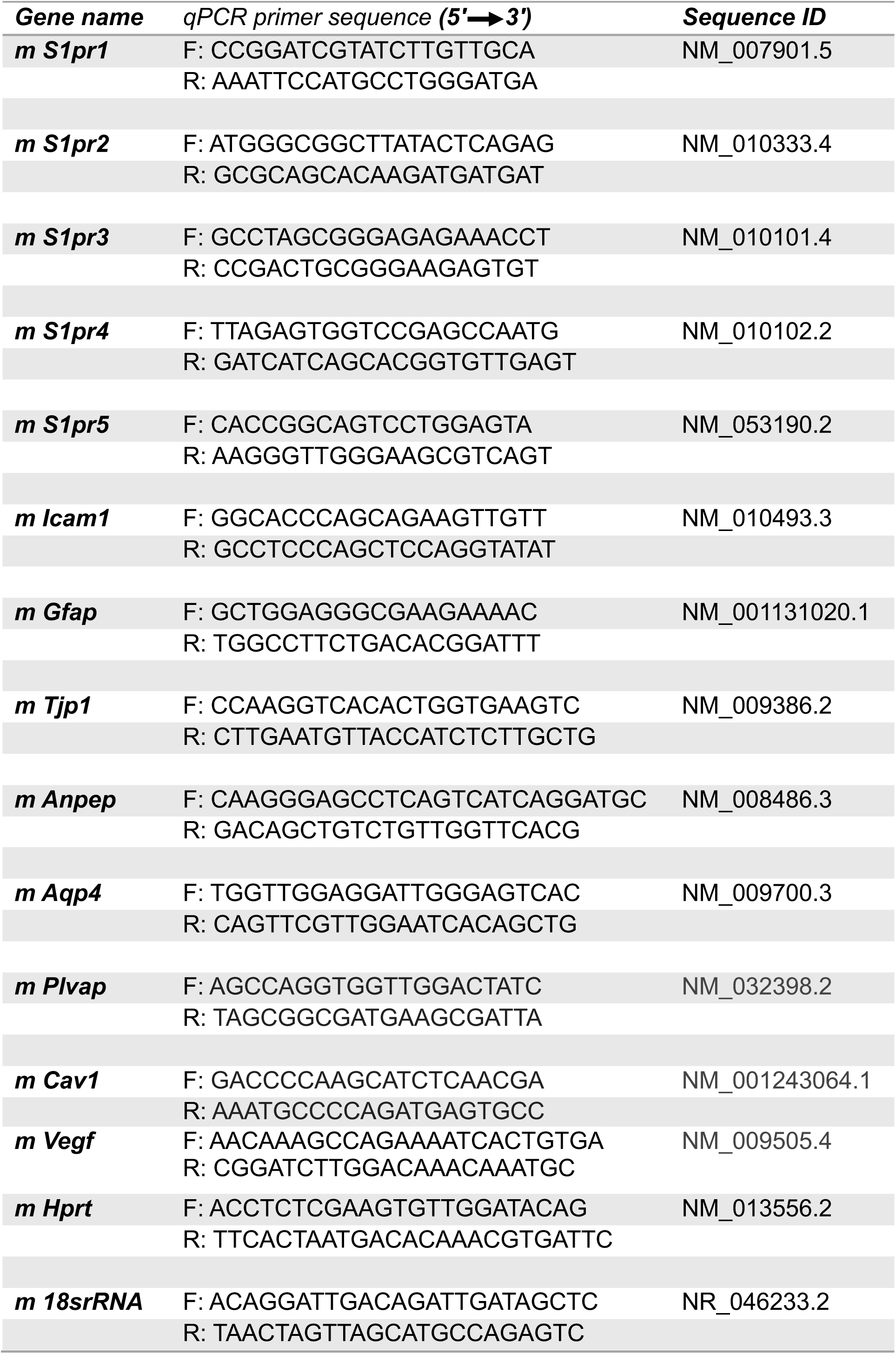
Sequences of the primers used for RT-qPCR.

#### S1PR1 Immunohistochemistry

Human tissues were retrieved from Brigham and Women’s Department of Pathology archives; this work was approved by the Institutional Review Board (Protocol #2013P001431). Immunohistochemistry for S1PR1 was performed on an automated stainer (Leica Bond III, Leica Biosystems, Buffalo Grove, IL) using an anti-human S1PR1 rabbit polyclonal antibody (Santa Cruz Biotechnology Inc.) at a final concentration of 1.3 ug/ml. The IHC technique for S1PR1 was validated as we have previously described ^62^. 5μm formalin fixed paraffin embedded tissue sections of human frontal cortex were deparaffinized and processed using heat induced epitope retrieval with an EDTA-based buffer (Leica #AR9640) for 20 minutes and incubated with primary antibody for 30 minutes at room temperature. Secondary antibody (polymer) incubation (10 minutes) and diaminobenzidine-based signal generation (10 minutes) were performed per manufacturer’s instructions (Leica # DS9800). Pictures were taken using SPOT Insight Gigabit camera and SPOT Imaging Software (5.1).

#### Immunofluorescence staining

Under deep anesthesia, mice were perfused with cold PBS and subsequently with 4% PFA in PBS solution. The brains were removed, postfixed with 4% PFA for 24 h, transferred to 30% sucrose solution in PBS, embedded in OCT media and frozen. Coronal sections were cut (9 µm) in a cryostat. Sections were washed three times with PBS and were then blocked with blocking solution (5 % bovine serum albumin, 0.8 % skim milk, and 0.3 % Triton X-100 in TBS) for 1 h and incubated with the specified primary antibodies in blocking solution overnight on a shaker at 4 °C, followed by the appropriate secondary antibodies and 4’,6-diamidino-2-phenylindole (DAPI) for 1 hours at room temperature and were mounted onto slides. Samples were observed on an FluoView FV10i confocal microscope (Olympus, Japan) (original magnification, x 60).

### Protein extraction from brain microvessels and western blotting

Brain microvascular fragments were lysed in HEPES-RIPA buffer (50 mM HEPES pH 7.5; 1% Triton; 0.5% sodium deoxycholate; 0.1% SDS; 500 mM NaCl; 10 mM MgCl2; 50 mM β-glycerophosphate) with 1x Protease inhibitor cocktail (CalBiochem), 1 mM Na3VO4 and 1 mM NaF and centrifuged at 15,000 r min^-1^ for 15 min. Equal amount of proteins were mixed with SDS sample buffer, boiled and separated on a 4-15% polyacrylamide gel (Bio-Rad), transferred to PVDF membranes (Millipore Sigma), and blocked 5 % milk in 0.1% Tween-20 in TBS. Immunoblot analysis was performed with S1PR1 (1:250; Santa Cruz, cat. no. sc25489, p-MLC (1:1000; Cell Signaling, cat. no. 3671), PLVAP (1:500; abcam, cat. no. ab27853), occluding (1:2000; Invitrogen, cat. no. 331500) and β-actin (1:1,000; Santa Cruz, cat. no. sc-1616 HRP) antibodies. Membranes were washed with 0.1% Tween-20 in TBS, incubated with anti-rabbit IgG secondary antibody conjugated to horseradish peroxidase (1:2,000; Cell Signaling), and protein bands were visualized with enhanced chemilumescent (ECL) reagent (Thermo Fisher Scientific) on a Protec OPTIMAX X-Ray Film Processor. Relative band intensities were obtained by densitometric analysis of images using ImageJ software.

#### Brain endothelial cell isolation and assessment of deletion efficiency of endothelial *S1pr1 mRNA*

Two weeks after tamoxifen treatment, mice were sacrificed and the brains were collected and rinsed in MCDB131 medium (Thermo Fisher Scientific) with 0.5% fatty acid-free BSA (Millipore Sigma). The cortices were homogenized in MCDB131 medium using a 7-mL loose-fit Dounce tissue grinder (Sigma-Aldrich) with 15 strokes. The homogenate was mixed with same amount of 30% dextran (molecular weight ∼70,000 Da, Sigma-Aldrich) in PBS and centrifuged at 4,500 r min^-1^ for 15 min at 4 °C. The pellet was resuspended in MCD131 medium and centrifuged at 2,400 r min^-1^ for 10 min. The pellet was resuspended in Liberase TM solution (3.5 U; Roche) with DNaseI (0.4 U; AppliChem Inc) and digested at 37.0 °C for 90 min. The enzymatic reaction was stopped by adding 2 mM of EDTA and 2% of BSA. After centrifugation at 2,400 r min^-1^ for 10 min, the pellet was incubated in purified Rat Anti-Mouse CD31antibody (MEC13.3) (1 : 100; BD Biosciences) with Dynabeads Sheep Anti-Rat IgG (Invitrogen, cat. no. 11035) for 35 min. CD31 positive endothelial cells were isolated by using DynaMag-2 Magnet (Thermo Fisher Scientific). Total RNA was extracted from the isolated endothelial cells using shredder (Qiagen) and RNeasy Mini Kit (Qiagen) with RNase-free DNase treatment (Qiagen) according to the manufacturer’s instructions. Reverse transcription was carried out using Verso cDNA Synthesis Kit (Thermo Fisher Scientific). Real-time PCR was performed on a real-time PCR system (Applied Biosystems, ABI 7500 Fast) by using PerfeCTa SYBR Green Fast Mix Low ROX. PCR primer sequences for target molecules are in Table 2.

#### Grading system for SAH

Blood volume in the subarachnoid space was assessed using the grading system previously reported ^63^. Mice were sacrificed under deep anesthesia 24 h after SAH induction and the brains were removed. Pictures of ventral surface of the brain depicting the basal cistern with the circle of Willis and the basilar artery were taken using a stereomicroscope (Olympus, SZX16) equipped with digital camera (Olympus, DP12). The basal cistern was divided into 6 segments and a grade from 0 to 3 was given to each segment: Grade 0, no subarachnoid blood; Grade 1, minimal subarachnoid blood; Grade 2, moderate blood clot with recognizable arteries; Grade 3, blood blot obliterating all arteries. Blood volume was evaluated by a total score ranging from 0 to 18 from six segments.

#### Tail bleeding assay

Tail bleeding time was determined as described previously^64^. A mouse was anesthetized with a mixture of ketamine and xylazine, and body weight was measured. The mouse was placed on a heating pad in prone position, the tail tip was about 4 cm blow the body horizon. A distal 5 mm segment of the tail was amputated, and the tail was immediately immersed in PBS pre-warmed at 37 °C. The time to complete arrest of bleeding was determined: complete arrest is no blood flow for 1 minute. The blood volume was determined by hemoglobin assay. Blood cells were separated by centrifuge at 4,000 r/min for 5 min at room temperature, and erythrocytes were resuspended in BD Pharm Lyse (BD Biosciences). After 10 min incubation in the buffer, the lysate was centrifuged at 10,000 rpm/min for 5 min. Hemoglobin concentrations were measured spectrophotometrically at 550 nm using a plate reader (Molecular Devices, SpectraMax M2e).

#### Mortality and neurological outcome

Mortality was assessed at 24, 48 and 72 hours after SAH induction. Gross neurological outcome was blindly evaluated before and at 24, 48 and 72 hours after surgery by sensorimotor scoring as described previously ^65, 66^. Briefly, a motor score (0 to 12; spontaneous activity, limb symmetry, climbing, and balance) and a sensory score were used. Gross neurological outcome was evaluated by a total score of 4 to 24. Higher scores indicate better neurological outcome.

#### Brain water content

Brain edema was determined by using the wet/dry method as previously described ^67^. Mice were sacrificed at 72 hours after surgery and the brains were quickly removed and separated into the left and right cerebral hemispheres and weighed (wet weight). The brain specimens were dried in an oven at 55°C for 72 hours and weighed again (dry weight). The percentage of water content was calculated as ([wet weight-dry weight]/wet weight) × 100%.

#### Cell death detection

DNA strand breakage during apoptosis after SAH was assessed by phospho-histone H2A.X (Ser 139) immunofluorescence as previously described ^68^. Because phosphorylation of histone H2A.X at Ser 139 (γ-H2AX) is abundant, fast, and correlates well with each DNA strand breakage, it is the most sensitive marker that can be used to examine the DNA damage ^69, 70^. Mice were deeply anesthetized and perfused with cold PBS and subsequently with 4% PFA in PBS solution. The brains were removed, post-fixed with 4% PFA for 24 h and transferred to 30% sucrose solution. Frozen brains were cut with a 10 µm of thickness by a cryostat (Leica, CM3050 S). The brain slices were blocked with TBS-blocking solution (5% bovine serum albumin and 0.5% Tween 20 in PBS) for 1 hour at room temperature and incubated with phospho-histone H2A.X (Ser 139) (20E3) antibody (1:100; Cell Signaling) in blocking solution overnight on a shaker at 4 °C. Sections were washed three times with 0.5% Tween 20 in PBS and then incubated with goat anti-rabbit IgG Alexa-488 (1:200; Life Technologies, cat. no. A-11008). DAPI (4’, 6’-diamidino-2-phenylindole) staining was used as a counter staining. Sections were imaged by using an FluoView FV10i confocal microscope (Olympus, Japan) (original magnification, x 40). For quantification, the percentage of phospho-histone H2A.X positive cells per DAPI positive cells from three different fields in the ipsilateral cerebral cortex area per mouse (bregma −1.64 to −1.28 mm) were counted and the average values were plotted.

#### Assessment of blood brain barrier dysfunction

To assess albumin extravasation, Evans blue dye (EBD) was used because EBD binds to plasma albumin^71^. Mice were anesthetized and 2% EBD (4 ml kg^-1^) was injected in the external jugular vein 21 hours after surgery. After 3 hours circulation, mice were deeply anesthetized and perfused with cold PBS to remove intravascular dye. The ipsilateral hemispheres were removed and homogenized in 50% trichloroacetic acid in PBS. The lysate was centrifuged at 15,000 rpm min^-1^ twice for 15 min at 4 °C, and the supernatant was used to measure fluorescence (excitation/ emission = 620/680 nm, SpectraMax M2e, Molecular Devices). The relative fluorescence units (R.F.U.) were normalized by the brain weights. To histologically confirm the plasma leakage into the brain parenchyma, 70 kDa dextran-TMR was injected through jugular vein and let circulate for 1h. Brain was removed without perfusion, embedded OCT compound directly and frozen ^72^. Sections were cut (9 µm) in a cryostat and fixed with 4% PFA before Immunohistochemistry. After staining with anti-Glut-1 antibody, the ipsilateral cortex was observed on an FluoView FV10i confocal microscope (Olympus, Japan) (original magnification, x 60).

#### Cell culture and treatments

Primary human brain endothelial cells HBMVECs (Cell systems, Kirkland, WA) were grown and maintained in 10cm dishes coated with attachment factor in complete media supplemented with culture boost serum and antibiotic bac-off as per manufacturer’s instructions. For *in vitro* assays, appropriate number of cells (Passage 4-8) were grown in culture plates to confluence. The brain endothelial cell line, bEnd.3 was obtained from ATCC (generous gift from Dr. Thomas White, Therapeutics Discovery Institute, Weill Cornell Medicine) and maintained in Dulbecco’s Modified Eagle Medium (DMEM) supplemented with 10 % fetal bovine serum (ATCC), and 100 U/ml penicillin/streptomycin (GIBCO) at 37 °C in a humidified 95 % air and 5 % CO2 incubator. Cells were pre-treated with 10µM of the S1PR1 antagonist, W146 (Cayman Chemical Co., USA), reconstituted in 100%Methanol (molecular grade) and or, 10µM Rho Kinase (ROCK) pharmacological inhibitor-Y-27632 (Cayman chemical Co., USA) reconstituted in distilled water, for 30min under appropriate conditions.

#### Genetic knockdown of mouse and human S1PR1

Genetic knockdown of human or mouse *S1pr1* mRNA was achieved using smartpool siRNA purchased from GE-Dharmacon (Horizon Discovery). 25nM of siRNA (human Cat No. L-003655-00-0005, mouse Cat No. L-051684-00-0005,) or scrambled control (D-001810-01-05) were transfected into HBMVECs or bEnd.3 cells (Passage 4-8) plated at a density of 150,000 cells per well in a 6-well culture plate using Lipofectamine 3000 (Life Technologies) as per manufacturer’s instructions in serum depleted and antibiotic free DMEM media. After transfection, cells were incubated for 36 h at 37°C/5% CO_2_, washed with PBS (1X), and harvested by gentle scraping for total RNA extraction.

#### Mechanistic assessment of brain endothelial responses to in vitro hypoxia-ischemia

To investigate the molecular and cellular mechanisms whereby S1PR1 signaling limits brain endothelial barrier dysfunction during SAH, HBMVEC or bEnd.3 cells (Passage 4-8) were used. Oxygen deprivation studies were conducted to recapitulate *in vitro* the ischemic-hypoxic environment of the cerebral microvasculature after SAH. S1PR1 signaling was blocked in HBMVEC or bEnd.3 cells by genetic knockdown (siRNA, 25nM) of S1PR1 or pharmacological inhibition (W146, 10μM). Cells were placed inside a portable hypoxic chamber (Coy Lab Products Inc., MI) while the chamber was purged with mixed gases (1%O_2,_ 95% Nitrogen, 4% CO_2_) for 4 minutes to maintain the inner chamber environment at depleted levels of oxygen (1%O2). Then the chamber with treated cells was placed in 37° C incubator for the times indicated. For normoxia condition, treated cells were incubated in normal culturing conditions.

#### Hypoxia-induced *in vitro* brain endothelial cell barrier dysfunction

HBMVEC cells (200,000 cells/insert) or bEnd.3 cells (100,000 cells/insert) were grown in complete growth media, as described above, in the Transwell Multiple Well Plate with Permeable Polyester Membrane Inserts (pore size 0.4 μm, culture area 0.33cm^2^, Costar, Corning, USA) coated with either attachment factor (Cell systems, Kirkland, WA) for HBMVECs or 50 μg/ml of human fibronectin (EMD Millipore) for bEnd.3. Upon confluence cells were treated with W146 and/or, Y27632 at 10µM in 0.4% fatty acid free BSA (Sigma Aldrich)/PBS (GIBCO) for 30m and then were placed inside a portable hypoxic chamber (Coy Lab Products Inc., MI) while the chamber was purged with mixed gases (95% Nitrogen, 4% CO_2_, 1%O_2_) for 4 minutes to maintain the inner chamber environment at depleted levels of oxygen (1%O2). HBMVEC cells were challenged with hypoxia window for 30min and bEnd.3 cells, for 6h, as determined from time course study. On the other hand, for normoxia condition, treated cells were incubated in normal culturing conditions for 6h or 30min respectively.

For genetic inhibition studies, cells were incubated with S1pr1 siRNA or scrambled control at 25nM for 24h at 37 degrees, 5%CO_2_. Cells were then treated with Y-27632 at 10µM for 30m at 37 degrees, 5%CO_2_ followed by incubation in hypoxic environment (1%O_2_) for an additional 6h inside the culture incubator. Thereafter, the chamber was opened to replenish normal atmospheric oxygen levels and 5 μl of 4.4KDa Tetramethylrhodamine Isothiocyanate (TMR)-Dextran (Sigma Aldrich) at 100mg/ml was added to each insert (upper chamber) and incubated for an additional 1h at 37 degrees, 5%CO_2_. The leakage of dextran was recorded by measuring the fluorescence intensity at Ex550nm/Em570nm with a fluorescence plate reader (SpectraMax M2, Molecular Devices) by taking 100 μl of medium from the lower chamber.

### Total RNA isolation, RT-qPCR analyses (*in vitro* studies)

Cells were rinsed with ice-cold PBS (1X) on ice twice after the treatments and TRI Reagent (Zymo Research) was added to each well and incubated for 3 min. The RNA was extracted from the lysate using Direct-zol RNA miniprep kit (Zymo Research) subjected to on-column DNaseI digestion according to the manufacturer’s instruction. Total RNA was eluted in nuclease free water and the concentrations were measured with multimode plate reader (Varioskan LUX, Thermo Scientific, USA). 100ng of total RNA was used to synthesize complementary DNA. Reverse transcription reaction was performed using Verso Reverse Transcriptase and Random Primer (Thermo Scientifc, USA) at 42 °C for 30 min. qPCR was performed using PerfeCTa SYBR Green FastMix Low Rox (Quantabio, USA) on ABI 7500 Sequence Detection System (Applied Biosystems, USA) with specific primer pairs (Table-2). The thermal cycling conditions for real time analysis were an initial denaturation at 95°C for 10 min followed by amplification and acquisition at 95°C for 10 sec, 56°C for 30 sec and 72°C for 30 sec for 30 cycles, and the thermal profile for melt curve analysis was obtained by holding the sample at 65°C for 31 sec followed by a linear ramp in temperature from 65°C to 95°C with a ramp rate of 0.5°C/sec and acquisition at 0.5°C intervals. The expression level of target mRNA obtained was normalized to that of *18srRNA* or *Hprt* mRNA expression. Relative expression of target mRNA in vehicle control and treated samples was expressed as 2-delta Ct values as described previously and fold changes were calculated by comparing the 2-delta Ct values of the treated sample with that of respective vehicle control.

#### Immunofluorescence imaging in HBMVEC and image analysis

HBMVEC were grown and stained in 35 mm glass bottom dishes (Mattek Corporation). In brief cells were fixed with 4% ice cold paraformaldehyde followed by permeabilization with 0.2% Triton-X 100. Cells were then blocked in 5% BSA in PBS at 4°C for 1h. Cells were then stained for CAV1 (Abcam, rabbit polyclonal, at 1:250), pMLC (Cell Signaling, rabbit polyclonal, at 1:100) or RAB5A (Abcam, mouse monoclonal, at 1:200). Appropriate goat anti-rabbit or donkey anti-mouse secondary antibodies conjugated with Alexa-fluor 488 or Alexa-fluor 594 was used for detecting the expression of CAV1, pMLC or RAB5A. For stress fiber detection, cells were incubated with fluorescent phalloidin (1:400 in PBS; Invitrogen) with emission at 549nm. Cells were then washed and stained with DAPI for nuclear staining. Images were acquired at 20X magnification using Biotek Lionheart FX microscope (Agilent) and analyzed using its built-in imaging software, Gen5. Images were then deconvoluted to remove any background. Changes in expression of total CAV1, pMLC and stress fibers were quantified using the automated imaging analysis algorithm using montage features at 20X magnification. The imaging montage parameters were set 300msec delay between channel movements followed by 8×8 grids in rows and columns. The horizontal spacing was set at 2894µM and vertical was set at 2791 µM from the center of the glass coverslip in clockwise manner with autofocus option Images were assigned to respective channels. For DAPI in the blue channel the LED intensity was set to 10 with integration time of 10msec and gain feature at 18 units. For CAV1 in green channel an LED intensity of 10 at integration time of 19msec and gain of 17units was optimized whereas, for pMLC in green channel an LED intensity of 10 at integration time of 38msec and gain of 150units was optimized given that the baseline for pMLC expression is relatively low. For endosomal marker, RAB5A in the red channel the LED intensity was set at 10 with an integration time of 540msec and camera gain of 25units. For phalloidin (stress fiber) stain visualization in the red channel an LED intensity of 10 with integration time of 19msec and gain of 15units was used.

For signal quantification, the mean intensity for total CAV1 or pMLC (in green) or Phalloidin (in red) channels was obtained for each 20X magnification image from the Gen5 image analysis software, BioTek Lionheart FX microscope. The total integral value of either green or red channel was normalized to the nuclei count in each image. Fold change in expression levels of the target proteins was expressed as relative to the scrambled siRNA or normoxia controls as presented in respective experimental analyses. Analysis was conducted from 7-10 independent images per experiment from a total of N=3-6 independent trials. For CAV1 and RAB5A punctates, similar analysis was performed with the spot count function in the Gen5 image analysis module to obtain the number of spots per cell per image.

#### CAV1 and RAB5A colocalization analysis

Co-localization analysis between CAV1 and RAB5A vesicle-like punctates was conducted as previously described ^73–75^ using Imaris version V9.9.1. Raw stack series images for each datapoint uploaded in the Imaris arena were first converted to the Imaris format (.ims) with x,y,z voxel parameters 0.4µM each. Images were assigned with respective channels from the color wheel. To identify punctates of CAV1 (green) and the early endosomal trafficking marker RAB5A (red), spots creation wizard was used. A minimum threshold was set for the spot size inclusion based on the XY diameter of spots and their pixel sizes that was identified as 0.6µM for red puncta and 0.65µM for green puncta. In-built machine learning algorithm was implemented to segregate the excluded spots. This step was included for both the green and red channels. Colocalized spots or punctates were identified by filtering the shortest distance between both the channels at a maximum threshold of 0.65µM, which is from the center of the CAV1 (green) punctates. This allows the detection of a population of red spots with centers located within 0.65µM from the center of CAV1 punctates, thus confirming a colocalization phenotype. The same algorithm was applied to quantify the number of CAV1 spots that overlapped with RAB5A punctates to confirm the phenotype.

#### Statistical analysis

All results were expressed as mean ± SEM. Statistical analysis were performed with GraphPad Prism (GraphPad Software, Version 7.0c) by using unpaired two-tailed Student’s t-test, one-way analysis of variance (ANOVA) or two-way ANOVA followed by Bonferroni’s or Tukey’s correction. *P* values < 0.05 were considered statistically significant.

## RESULTS AND DISCUSSION

### Endothelial S1PR1 is an endogenous protective pathway which limits SAH-induced cerebral edema

We first investigated the expression of S1PR1 in the mouse brain using the S1pr1-eGFP knock in mice, in which S1PR1-eGFP fusion protein is expressed under the control of the endogenous S1PR1 promoter ^58^. Consistent with our recent RNA analyses by *in situ* hybridization approach ^76^, we found that S1PR1-eGFP protein is abundantly expressed in the endothelium of cerebral microvessels throughout the brain (Figure 1A-C and Supplemental Figure 1) as assessed by detection of S1PR1-eGFP and immunofluorescence (IF) analysis for the glucose transporter 1 (Glut-1). S1PR1 is also expressed in parenchymal cells throughout the brain, mainly in the neuropil, as assessed by microtubule associated protein (MAP)-2 and Nissl staining (Supplemental Figures 2, 3). Only some fibrous astrocytes were found to express S1PR1 (Supplemental figure 4). Next, we quantified the expression of S1PR1 mRNA by reverse transcription and quantitative PCR analysis (RT-qPCR), as we have previously described ^62, 77^ and compared it to the expression of the other S1PR isotypes. We found that S1PR1 is the most abundant S1PR transcript in the mouse brain (20.5± 2.4 copies/10^6^ 18S, which is equivalent to approximately 20.5± 2.4 S1PR1 mRNA copies/cell ^61^, Supplemental figure 5A). Also, comparison of the S1PR1 mRNA levels in cerebral microvessels with the brain parenchyma, revealed that S1PR1 transcripts are highly enriched in cortical microvessels when compared to total brain (Supplemental figure 5B, ∼6-fold enrichment in microvessels *versus* whole brain). Altogether, these data indicate that S1PR1 is the most abundant S1PR transcript in the brain and is highly enriched in mouse cerebral microvessels when compared to total brain. At the protein level, S1PR1 is also widely expressed in the cerebrovascular endothelium in the mouse brain and the grey matter neuropil.

**Figure 1.**
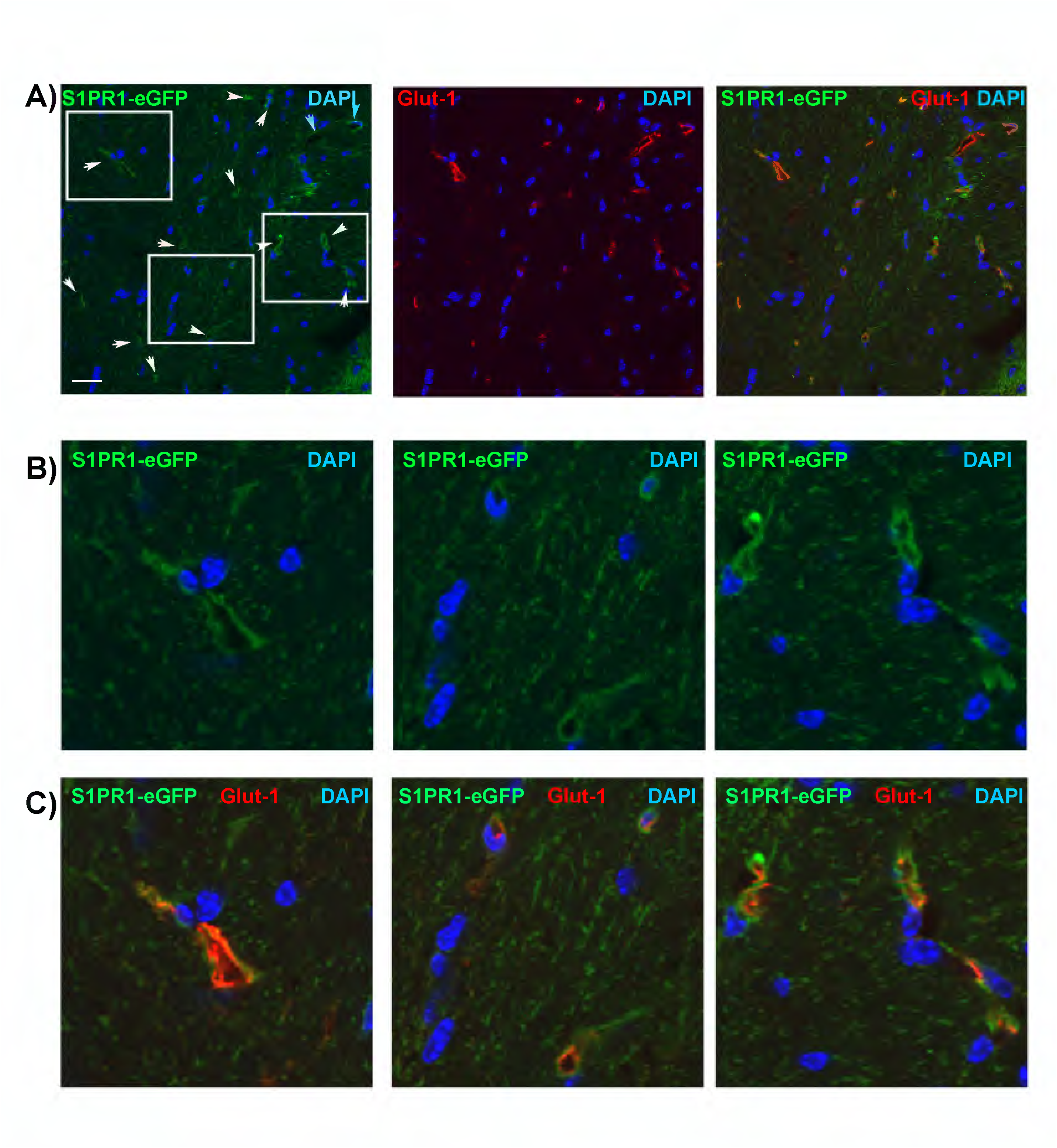
S1PR1 is expressed in the cerebrovascular endothelium of the mouse brain. Confocal analysis of S1PR1-eGFP fluorescence and Glut-1 immunodetection of the mouse brain. A) S1PR1-eGFP (green channel) is expressed in the cerebrovascular endothelium. Immunofluorescence for Glut-1 (endothelial marker) is shown in the red channel. Representative image of the corpus callosum. Scale bar 20μm. N=5-6. Sections were captured using an FluoView FV10i confocal microscope (Olympus, Japan) (original magnification, x 60). Notice the high expression of S1PR1-eGFP in microvessels (white arrows). B-C) High power view (digital zoom) of selected areas in panel A.

In order to investigate the role of endothelial S1PR1 signaling in the pathophysiology of early brain injury after aneurysmal SAH, we used the endovascular rupture model of SAH ^60^, which is a well-established and reproducible model that recapitulates key features of the pathophysiology of the acute phase of SAH, such as an increase in intracranial pressure when blood pours/flows into the subarachnoid space upon rupture of the MCA, which gives rise to a brief period of transient (∼3-5 minutes) cerebral ischemia ^60^, followed by gradual recovery of the CBF and hypoperfusion in the acute phase, cerebral edema and neuronal injury ^16, 19^. Wild-type mice were subjected to SAH as we described and 24 hours later, cerebral microvessels were isolated from the mouse cortex to determine S1PR1 mRNA and protein levels compared to sham mice. We found that S1pr1 mRNA levels in cortical microvessels were significantly increased at 24h after SAH (3.14±0.55-fold induction) compared to sham (Figure 2A), together with the levels of the markers of neurovascular inflammation intercellular adhesion molecule (Icam)-1 and Gfap mRNA (Figure 2B and 2C, 3.6±0.37 and 5.14± 0.55-fold induction, respectively). In contrast, the levels of endothelial (tight junction protein-1, Tjp-1, Figure 2D), pericyte (CD13, Anpep, Figure 2E) or astrocyte end foot (Aqp4, Figure 2F) markers did not significantly change suggesting similar cell composition among the microvessel preparations from these groups of mice. Western blot analysis of cerebral microvessels confirmed the increase in S1PR1 protein levels at 24h after SAH compared to sham (1.68 ± 0.19-fold, Supplemental Figure 6). These data indicate that S1PR1 mRNA and protein levels in cerebral microvessels are significantly increased during the acute phase of SAH.

**Figure 2.**
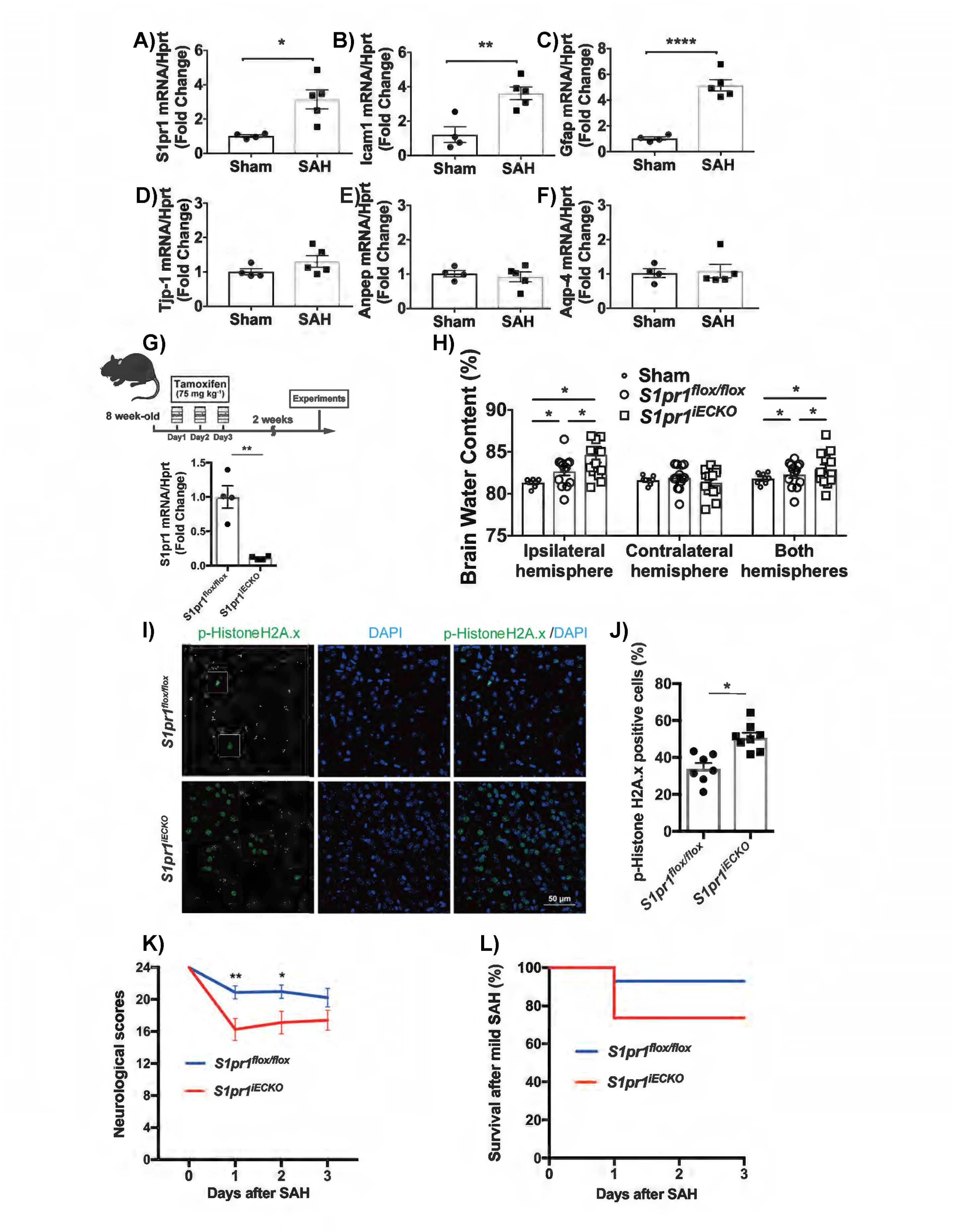
Endothelial S1PR1 is an endogenous protective pathway that limits cerebral edema and neuronal injury after SAH. A-F) Alterations in S1pr1, Icam-1, Gfap, Tjp-1 (ZO-1), Anpep (CD13) and Aqp4 mRNA levels in cerebral microvessels were quantified by RT-qPCR in sham animals 24 h after SAH. S1pr1, Icam-1 and Gfap are induced in cerebral microvessels after SAH. The mRNA levels of the endothelial (ZO-1, Tjp-1), pericyte (Anpep, CD13) or astrocyte (Aqp4) markers did not significantly change suggesting similar cell composition among the microvessel preparations from these groups of mice. G) Scheme of tamoxifen treatments and efficiency of deletion of S1PR1 in the cerebrovascular endothelium. *S1pr1^flox/flox^* and *S1pr1^iECKO^*mice were treated with tamoxifen (75 mg kg^-1^) for the consecutive 3 days at the age of 8 weeks. *S1pr1* mRNA levels were analyzed by qPCR in brain endothelial cells isolated from *S1pr1^flox/flox^*and *S1pr1^iECKO^* mice, 2 weeks after tamoxifen treatment (*n* = 4). Efficiency of deletion in the cerebrovascular endothelium in *S1pr1^iECKO^* mice is shown relative to *S1pr1^flox/flox^*mice. Individual values and mean ± SEM are shown. \**P* < 0.05, \*\**P* < 0.01, \*\*\**P* < 0.001, Student’s t-test. H) 72 hours after SAH or sham surgery, brain edema was evaluated by quantification of brain water content in *S1pr1^flox/flox^* and *S1pr1^iECKO^* mice (*n* = 7 - 14). Brain water content in the ipsilateral and contralateral hemispheres was calculated as ([wet weight-dry weight]/wet weight) × 100. * p<0.05, one-way ANOVA followed by Tukey’s test. I) Immunofluorescence confocal analysis of phospho-histone H2A.X, a marker for DNA damage (green channel). Nuclear staining (DAPI) is shown in the blue channel. Representative images of ipsilateral hemisphere (cortex) from *S1pr1^flox/flox^* and *S1pr1^iECKO^* mice after SAH stained are shown. Scale bar, 50 µm. J) Quantification of phospho-histone H2A.X positive cells (%) (n = 7-8 mice). \**P* < 0.05, student’s t-test. The individual values and the mean ± SEM are shown. Each data point represents a mouse and it is the average of 3 different fields (bregma −1.64 to −1.28 mm). K) Neurological (sensory and motor) deficits after SAH surgery in *S1pr1^flox/flox^* and *S1pr1^iECKO^* mice (n = 12 or 14) were assessed as described in methods. From 4 to 24 points: 24 points (best), 4 points (worst). Data are mean ± SEM. \**P* < 0.05, \*\**P* < 0.01, One-way ANOVA followed by Tukey’s test. Red line, *S1pr1^iECKO^*; Blue line, *S1pr1^flox/flox^*. L) Survival curves in *S1pr1^flox/flox^*and *S1pr1^iECKO^* mice after SAH surgery (*n* = 13 or 19). There were not statistically significant differences in survival by Log-rank test. Red line, *S1pr1^iECKO^*; blue line, *S1pr1^flox/flox^*.

Previous studies indicated that inhibition of S1PR1 signaling via administration of the immune suppressor FTY720 protected in experimental subarachnoid hemorrhage ^50, 51^ possibly via their immunosuppressive effects ^48^ ^30^. Given the abundant expression of S1PR1 in the cerebrovascular endothelium, its enrichment in cerebral microvessels, and its induction by SAH, we aimed to investigate the role of endothelial S1PR1 signaling in early brain injury after SAH. We generated endothelial-specific S1PR1 null mice (*S1pr1^iECKO^*) by treating adult *S1pr1^flox/flox^xCdh5–Cre^ERT2^* mice (∼2-month-old) with 75 mg kg^-1^ tamoxifen for 3 consecutive days (Figure 2G) as we have recently described ^55^. *S1pr1^flox/flox^* littermate mice were also treated with tamoxifen and used as wild-type control. RT-qPCR analysis of isolated brain endothelial cells 2 weeks after tamoxifen treatment demonstrated a ∼90% reduction in S1pr1 expression in S1pr1iECKO mice compared to S1pr1flox/flox littermate mice (Figure 2G). *S1pr1^iECKO^* and *S1pr1^flox/flox^* littermates were subjected to SAH surgery using a modified 5.0 suture (0.3 mm x 0.3 mm, as described in methods section). To determine the impact of the lack of endothelial S1PR1 signaling on brain injury after SAH, we assessed brain edema 72 hours after the induction of SAH by quantifying total brain water content. Consistent with previous findings using the rupture model of SAH ^11, 60^, we observed a modest but statistically significant increase in total brain water content in the ipsilateral hemisphere 72 h after SAH in *S1pr1^flox/flox^*mice compared to sham (82.83 ± 0.53% compared to 81.34 ± 0.20%, Figure 2H, 1.8% increase *vs* sham). Interestingly, in S1pr1iECKO mice, total brain edema in the ipsilateral hemisphere after SAH was significantly higher than that in *S1pr1^flox/flox^* (84.39 ± 0.91% compared to 82.83 ± 0.53%, Fig. 2H), a 3.75% increase compared to sham. There were no significant changes in the brain water content in the contralateral hemisphere (SAH *vs* sham) in either *S1pr1^flox/flox^ or S1pr1^iECKO^* mice.

Next, we analyzed cell death at 24 hours after SAH using phospho-histone H2A.X (Ser 139) immunostaining, a marker for double strand DNA breaks and neuronal endangerment ^69, 70, 78, 79^. We found that *S1pr1^iECKO^* mice showed significantly higher numbers of phospho-histone H2A.X positive cells compared to *S1pr1^flox/flox^*mice (50.85 ± 2.56% in *S1pr1^iECKO^* versus 34.00 ± 2.98% in *S1pr1^flox/flox^*) (Figure 2I, J). No phospho-histone H2A.X positive cells were detected in sham animals. Neurological outcomes were determined by assessing motor and sensory function using a total scale of 4 to 24 (being 24 the best neurological outcome) as previously described^65, 66^. Neurological outcomes at 24h and 48h after SAH surgery were worse in *S1pr1^iECKO^* mice (16.29 ± 1.37 and 17.14 ± 1.40, respectively) than in *S1pr1^flox/flox^* mice (20.91 ± 0.81 and 21.00 ± 0.83, respectively) (Figure 2K). Mortality rates on days 1, 2, and 3 after SAH were 7.7% in wild-type mice (*S1pr1^flox/flox^*) and 26.3% in S1pr1iECKO mice; however, the differences in mortality were not statistically significant (Figure 2L).

These data indicate that genetic deletion of *S1pr1* specifically in the endothelium significantly exacerbates total brain edema and neuronal injury resulting in poorer neurological outcomes.

### Genetic deletion of S1pr1 in the endothelium does not affect the key vascular and physiological parameters regulating blood volume in the subarachnoid space or cerebral blood flow changes

The volume of blood in the subarachnoid space, which depends on the amount of bleeding and clearance, is directly correlated with worse outcomes after SAH ^80^ ^63^. Thus, we quantified the amount of subarachnoid blood upon SAH in *S1pr1^flox/flox^* and *S1pr1^iECKO^* mice by image analysis using a previously described SAH grading system ^63^. No significant differences were observed in the amount of subarachnoid blood between *S1pr1^flox/flox^* and *S1pr1^iECKO^*mice at 24h after SAH (Fig. 3A, representative pictures). SAH grading was 11.50 ± 0.62 in *S1pr1^flox/flox^* mice and 11.67 ± 0.42 in *S1pr1^iECKO^* mice (Fig.3B). In addition, we determined the role of endothelial-specific S1PR1 in hemostasis using the tail bleeding assay ^64^. We did not find any significant differences in bleeding times or blood volumes between *S1pr1^flox/flox^* and *S1pr1^iECKO^* mice in the tail bleeding assay. Bleeding times were 68.88 ± 4.98 seconds in *S1pr1^flox/flox^* and 63.8 ± 4.14 seconds in *S1pr1^iECKO^* (Fig. 3C). Hemoglobin content, assessed by measuring the absorbance at 550 nm, was 0.73 ± 0.07 in *S1pr1^iECKO^* mice compared to 0.70 ± 0.12 in *S1pr1^flox/flox^* mice (Fig. 3D). Altogether, these data indicate that the amount of blood in the subarachnoid space 24 h after SAH is similar in wild-type and *S1pr1^iECKO^ mice and* that hemostasis is not altered in *S1pr1^iECKO^* mice compared to wild-type mice. Thus, the worsened SAH outcomes observed in *S1pr1^iECKO^* mice compared to wild type cannot be explained by differences in bleeding or clearance of the subarachnoid blood upon endovascular rupture.

**Figure 3.**
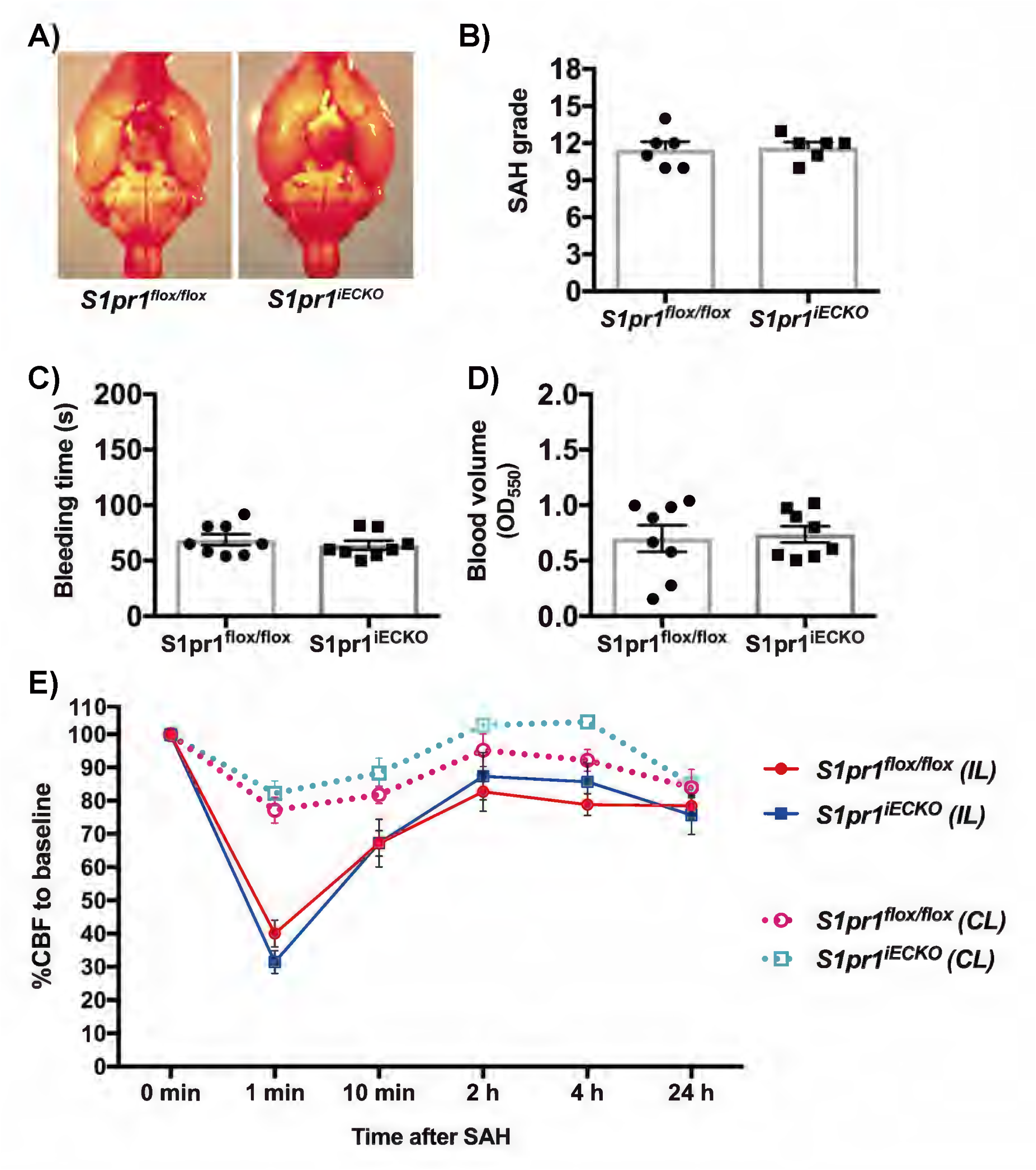
Genetic deletion of *S1pr1* in the endothelium has no effect on subarachnoid blood volume, hemostasis or CBF changes after SAH. A) Representative images of the ventral side of the brain from *S1pr1^flox/flox^* and *S1pr1^iECKO^* mice at 24 hours after SAH. B) Subarachnoid blood volume (SAH grade) was calculated by quantifying the blood in the subarachnoid space by image analysis as described in methods section. (*n* = 6). (C-D) Quantification of hemostasis in the tail bleeding assay. C) bleeding time and D) blood volume (*n* = 8). (B-D) Student’s t-test. Individual values and mean±SEM are shown. (E) CBF in the middle cerebral artery territory was measured by Laser-speckle contrast imager in *S1pr1^flox/flox^* and *S1pr1^iECKO^* mice before (0 min.), 1 min, 10 min, 2 h, 4 h, 24 h after SAH induction. The relative CBF values (%) *versus* before SAH are shown. Data are mean ± SEM. (*n* = 6). Two-way ANOVA followed by Bonferoni’s test showed no statistically significant differences between wild type (*S1pr1^flox/flox^*) and *S1PR1^iECKO^*. Red solid line, *S1PR1^flox/flox^* ipsilateral hemisphere (SAH affected side); blue solid, *S1PR1^iECKO^* ipsilateral; pink dotted line, *S1PR1^flox/flox^* contralateral; light blue dotted line, *S1PR1^iECKO^* contralateral. IL, ipsilateral side. CL, contralateral side.

We also determined the physiological parameters that could have an impact on CBF changes after SAH. No significant differences were observed in arterial O_2_ saturation, heart rate, pulse distention (a surrogate of pulse pressure), and respiratory rate between *S1pr1^iECKO^*and *S1pr1^flox/flox^* mice, before or after SAH (Table 1). Next, we quantified the cerebral blood flow (CBF) changes during SAH in both groups of mice using laser speckle flowmetry (Figure 3E). CBF in the middle cerebral artery (MCA) territory rapidly dropped ∼1 minute after SAH in a similar manner in both *S1pr1^flox/flox^* and *S1pr1^iECKO^* mice. In the ipsilateral hemisphere, CBF dropped to 40.26 ± 4.09% of basal in *S1pr1^flox/flox^ and* to 31.65 ± 3.46% of basal in *S1pr1^iECKO^*; in the contralateral hemisphere, CBF dropped to 77.32 ± 3.90% of basal in *S1pr1^flox/flox^* and to 82.42 ± 3.72% of basal in *S1pr1^iECKO^*. Afterwards, as shown in Figure 3E, CBF progressively recovered similarly in S*1pr1^flox/flox^* and *S1pr1^iECKO^* mice. 2 h after SAH, in the ipsilateral hemisphere, CBF recovered to 82.83 ± 5.88% of basal in *S1pr1^flox/flox^ and* to 87.46 ± 7.15% of basal in *S1pr1^iECKO^*; in the contralateral side CBF recovered to 95.34 ± 4.86% of basal in *S1pr1^flox/flox^* and to 102.81 ± 1.97% of basal in *S1pr1^iECKO^*. At 4h and 24h after SAH, the ipsilateral hemisphere remained hypoperfused (CBF was ∼20% lower than basal) in both S*1pr1^flox/flox^*and *S1pr1^iECKO^* mice. These data indicate that genetic deletion of *S1pr1* in the endothelium does not have a significant impact on CBF changes upon SAH.

Altogether, these data indicate that endothelial-specific deletion of S1PR1 in adult mice does not have a significant impact on bleeding or clearance of subarachnoid blood, or systemic physiological parameters such as heart rate, respiratory rate, O_2_ saturation, pulse distension or cerebral blood flow changes upon SAH.

### Endothelial-specific deletion of *S1pr1* in adult mice exacerbates blood brain barrier dysfunction following SAH

Since endothelial S1PR1 deletion did not alter physiological and systemic vascular parameters that could impact blood accumulation in the subarachnoid space, and CBF changes after SAH, we focused our investigation on the cerebral microvasculature and set out to determine blood-brain barrier permeability after SAH in wild-type and *S1pr1^iECKO^*mice. First, we quantified albumin leakage using the Evans Blue dye (EBD) extravasation assay. 24 h after SAH, albumin leakage in the ipsilateral hemisphere was significantly increased in *S1pr1^flox/flox^* mice compared to sham animals (Figure 4A, 1.53 ± 0.06-fold in *S1pr1^flox/flox^* SAH versus *S1pr1^flox/flox^*sham). Remarkably, SAH-induced BBB permeability was significantly higher in *S1pr1^iECKO^* mice than that in *S1pr1^flox/flox^* (1.83 ± 0.08 in *S1pr1^iECKO^*, Figure 4A). No differences in albumin leakage were observed between *S1pr1^flox/flox^* and *S1pr1^iECKO^* mice subjected to sham surgery, which is consistent with our recent report ^55^. We also aimed to determine the effect of sex and age on the phenotype observed in *S1pr1^iECKO^*. As shown in Supplemental Figure 7A, like the phenotype observed in male mice, female *S1pr1^iECKO^* exhibited exacerbated BBB leakage compared to wild type. Mortality rates were also similar in female *S1pr1^iECKO^* (10%) mice compared with wild-type mice (8.3%). In middle-aged mice (46-52 weeks old), mortality at 24 h after SAH was higher than that in young mice but similar between the wild-type (48%) and *S1pr1^iECKO^* (39%) groups. BBB permeability was significantly higher in aged *S1pr1^iECKO^* mice compared to *S1pr1^flox/flox^ mice* (Supplemental Figure 7B). Thus, endothelial S1PR1 limits BBB permeability after SAH both in males and females and in aged mice.

**Figure 4.**
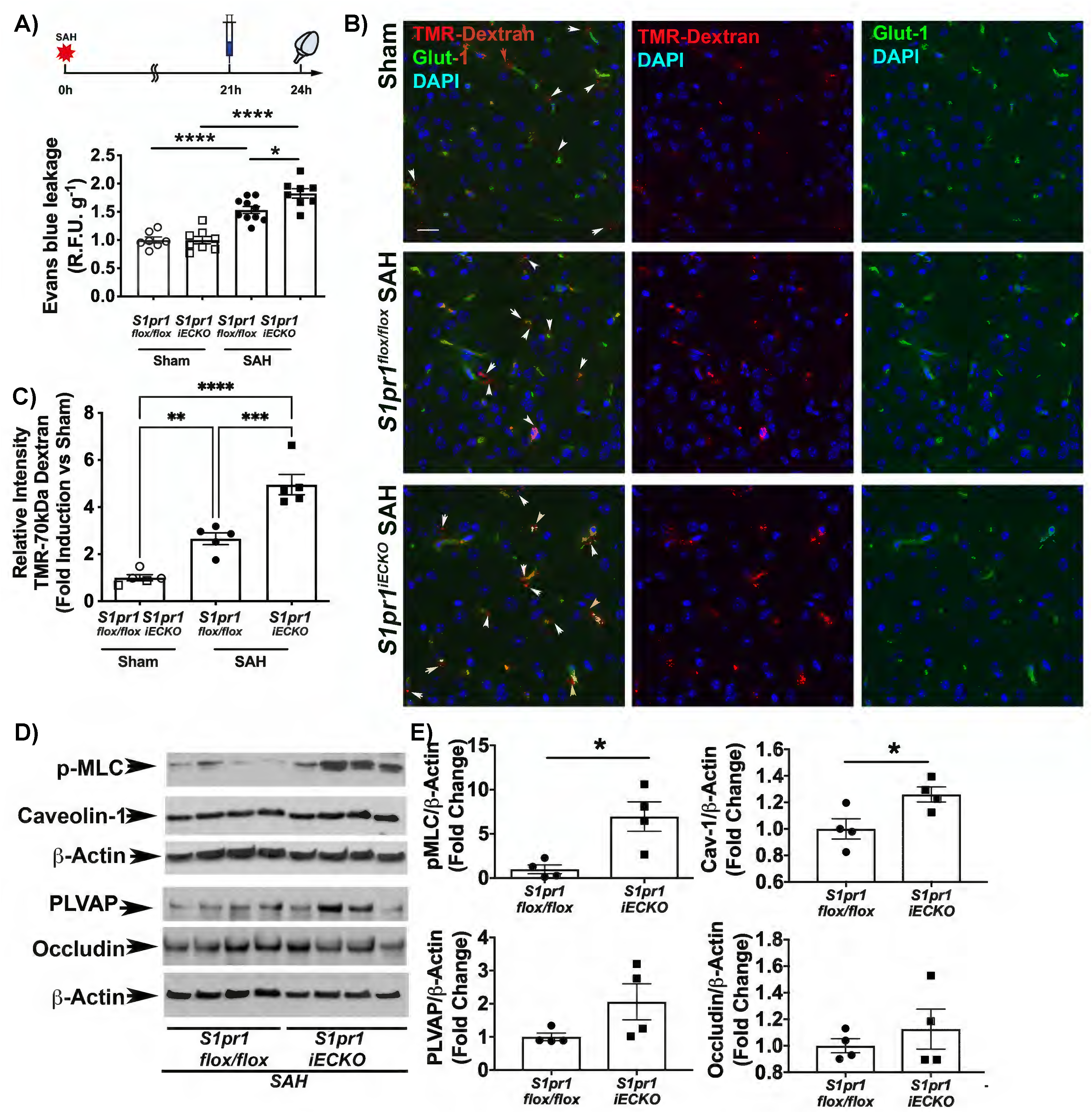
Endothelial specific S1PR1 null mice exhibit exacerbated BBB leakage and increased expression of molecular markers of barrier dysfunction in cerebral microvessels after SAH compared to wild type littermates. A) Albumin BBB leakage, assessed by Evans Blue Dye extravasation, 24 hours after sham or SAH surgery in *S1pr1^flox/flox^* and *S1pr1^iECKO^* mice (*n* = 6 - 10). Evans blue dye was circulating for 3 hours. The individual values and mean ± SEM are shown. \**P* < 0.05, \*\**P* < 0.01, \*\*\**P* < 0.001, \*\*\*\**P* < 0.0001 (One-way ANOVA followed by Tukey’s test). *y* axis shows relative fluorescence units (R.F.U.) per gram of tissue normalized *vs* sham. B) Histological confirmation of BBB leakage. 70kDa TMR-dextran was used as tracer. Immunofluorescence for Glut-1 (endothelial marker) is shown in the green channel. Nuclear stain (DAPI) is shown in the blue channel. White arrows point to extravascular (parenchymal) TMR-dextran. Representative pictures from cortex are shown. Scale bar 20μm. N=5-6. Sections were captured using an FluoView FV10i confocal microscope and a 60x objective (Olympus, Japan). C) Quantification of 70kDa of TMR-dextran leakage into the brain parenchyma. D-E) Increased levels of phosphorylated myosin light chain (p-MLC) and caveolin 1 in isolated cortical microvessels in *S1pr1^iECKO^* mice compared to *S1pr1^flox/flox^*. Cortical microvessels were isolated 24 h after SAH in *S1pr1^iECKO^* and *S1pr1^flox/flox^* mice. D) Western blot analysis for phospho-MLC (p-MLC), caveolin-1, PLVAP, occludin and β-actin. E) Western blot quantification was conducted using Image J. Individual values and mean ± SEM are shown. \**P* < 0.05 (t test).

Next, we histologically confirmed the leakage of macromolecules into the brain parenchyma by intravenous injection of 70kDa tetramethylrhodamine (TMR)-dextran and immunofluorescence analysis of the endothelial marker Glut-1. Histological analysis of 70kDa tetramethylrhodamine (TMR)-dextran localization, confirmed the leakage of the intravascular tracer into the brain parenchyma and outside of the cerebral capillaries 24 h after SAH (Figure 4B, C). Quantification of extravascular TMR-dextran confirmed that SAH-induced BBB permeability was higher in *S1pr1^iECKO^* mice than in *S1pr1^flox/flox^*(4.9±0.43 vs 2.6±0.25, respectively). Altogether, these data indicate that BBB dysfunction upon SAH is exacerbated in both male and female young and middle-age mice lacking S1PR1 specifically in the endothelium, compared to that in wild-type.

To elucidate the mechanisms by which endothelial deletion of S1PR1 leads to exacerbated BBB dysfunction, we isolated cerebral microvessels after SAH from *S1pr1^flox/flox^ S1pr1^iECKO^* mice and determined the activation of signaling pathways implicated in BBB dysfunction by western blot analysis. The Rho-ROCK pathway has been shown to play a critical role in endothelial dysfunction and inflammation ^81^ in peripheral organs, as well as blood-brain barrier dysfunction in numerous CNS pathological conditions ^82–85^. Rho-ROCK promotes the formation of actin stress fibers and assembly of the actomyosin apparatus, via phosphorylation and activation of MLC, leading to endothelial contraction and BBB leakage^82^. We quantified the levels of phosphorylated myosin light chain (MLC), a downstream effector of the Rho-ROCK pathway critical for endothelial cytoskeleton rearrangements (e.g. actomyosin assembly and cell contraction) leading to blood-brain barrier dysfunction ^82^. We found that exacerbated BBB dysfunction in endothelial-specific S1PR1 null mice was associated with increased phosphorylation of myosin light chain (MLC) in isolated cortical microvessels (Figure 4D, E, 6.96±1.6 fold in *S1pr1^iECKO^ versus S1pr1^flox/flox^*). We also observed that the protein levels of Caveolin-1 (Cav-1), which plays a key role in vesicular trafficking and endothelial transcytosis^86^, were modestly but significantly increased in *S1pr1^iECKO^* mice compared to *S1pr1^flox/flox^* (1.26±0.056 fold increase, Figure 4D, E). A marked trend towards increased levels of the caveolar protein PLVAP (1.8±0.38-fold, p=0.06, Fig. 5D, E). In contrast, the levels of the tight junction protein occludin were similar in the wild-type and *S1pr1^iECKO^ mice (*Figure 4D, E) ^59^.

**Figure 5.**
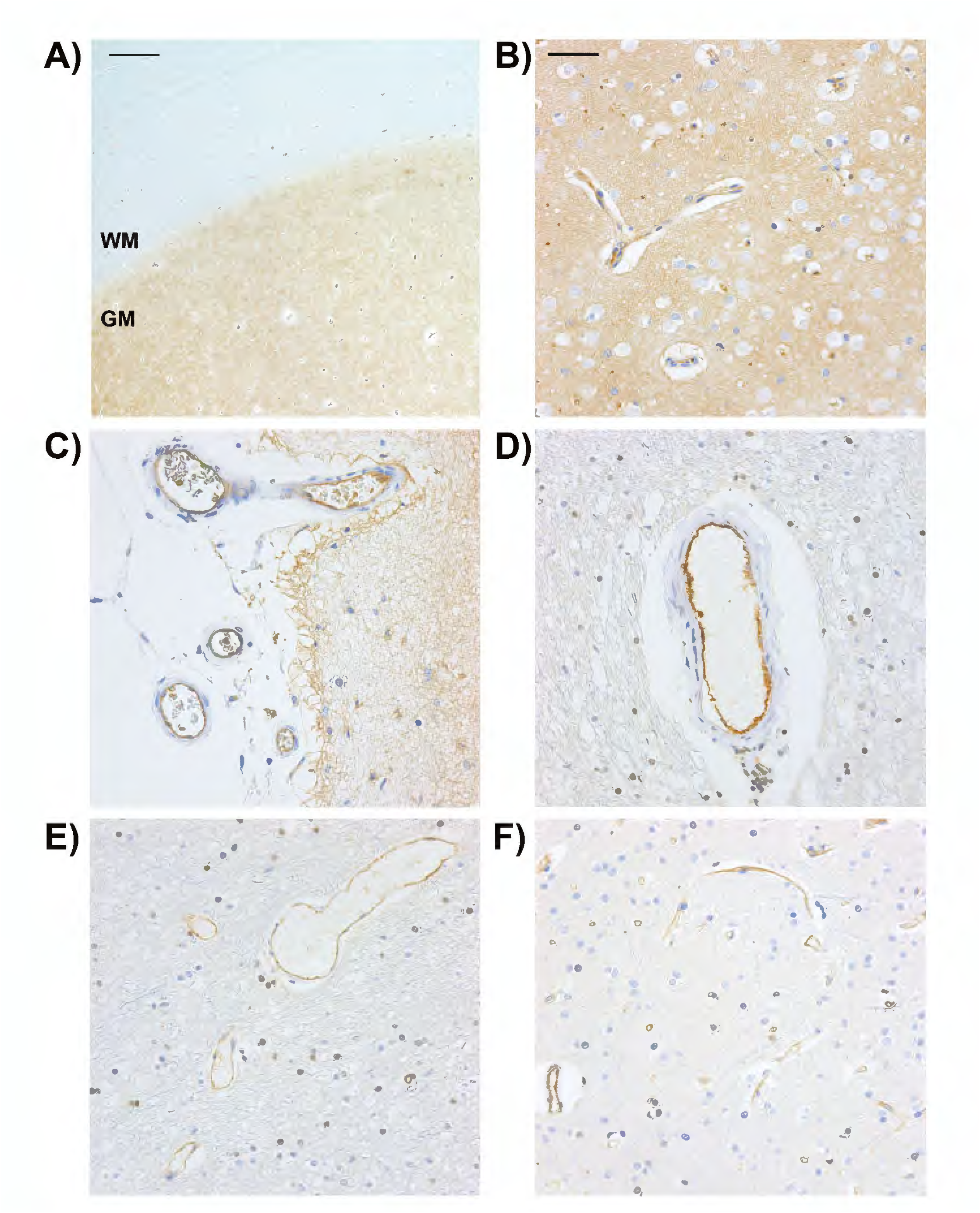
Detection of S1PR1 in the human brain. Representative images of S1PR1 immunohistochemistry from 5 human brain autopsy samples. Formalin-fixed paraffin-embedded tissue sections were used for immunohistochemical staining of S1PR1, as described in the Methods section. A) Low magnification picture showing the detection of S1PR1 in frontal cortex grey matter (GM) and subcortical white matter (WM). Scale bar 500 μM. B) Representative picture showing S1PR1 immunopositivity in arterioles and capillaries as well as the neuropil of the cortical grey matter. C) Detection of S1PR1 in the cerebrovascular endothelium of pial vessels. D-F) Representative pictures of S1PR1 immunodetection in the subcortical white matter. Notice S1PR1 positivity in parenchymal arterioles (D, F), venules (E) and capillaries (E, F). B-F) Scale bar 50 μM. Pictures were taken with a 60x objective (Olympus, Japan).

These data indicate that BBB dysfunction (albumin leakage) after SAH is heightened in mice lacking S1PR1 specifically in the endothelium, which is associated with increased levels of phosphorylation of the Rho-ROCK downstream effector, MLC, and caveolar proteins involved in endothelial transcytosis in cerebral microvessels. Altogether, our data point to endothelial S1PR1 as an endogenous protective mechanism to limit SAH-induced BBB dysfunction and overactivation of the pro-inflammatory Rho-ROCK pathway.

### S1PR1 is widely detected in the human cerebrovascular endothelium

To investigate the expression of S1PR1 in the human brain, we performed IHC analysis. The S1PR1 antibody and IHC protocol used for human samples have been previously validated in our laboratory ^62^. Representative images of the frontal cortex are shown in Figure 2. S1PR1 immunostaining was observed in both grey matter (GM) and subcortical white matter (WM) areas (Figure 5A, low magnification). S1PR1 was widely detected in the cerebrovascular endothelium of parenchymal vessels (in the grey matter and white matter, Figure 5B, D-F) and pial vessels (Figure 5C). In the grey matter, S1PR1 staining was also observed in the neuropil (Figure 5B, C). In the subcortical white matter, the signal intensity of S1PR1 was more scattered and weaker compared to the strong signal observed in the microvessels (Figure 5E and F). These data indicate that S1PR1 is prevalent in the cerebrovascular endothelium of humans.

### S1PR1 is protective signaling pathway in human cerebral microvascular endothelial cells that restrains hypoxia-induced barrier leakage, MLC phosphorylation, stress fiber assembly and Cav-1 distribution to endosomes

Given the key role of S1PR1 in limiting BBB leakage in mice and its abundant detection in the human cerebrovascular endothelium, we set out to investigate the functional role of S1PR1 in barrier permeability in primary human brain microvascular endothelial cells (HBMVEC). We used an *in vitro* system to recapitulate the hypoxic microenvironment of the cerebrovascular endothelium during SAH^7^, ^8^, as we have previously described^87^. Primary human brain microvascular endothelial cells (HBMVEC) were cultured in Transwell plates and subjected to hypoxia (1% O2) to induce barrier dysfunction. Barrier permeability was assessed by tracer leakage (TRITC-Dextran) from the top to the bottom chamber. We initially determined the severity of the hypoxic challenge that induced robust barrier leakage by subjecting the cells to different times of hypoxia. In HBMVEC, a robust increase in barrier leakage was induced after 30 min of hypoxic challenge (1.8±0.15 fold-induction hypoxia vs normoxia, Supplemental figure 8A and C). Hypoxia-induced barrier dysfunction was associated with a strong induction of VEGF and Cav-1 mRNA in HBMVEC (9.1±1.8 fold and 3.1±0.3 fold, respectively) (Supplemental Figure 8E, F). To determine the role of S1PR1 in hypoxia-induced barrier dysfunction, HBMVEC were transfected with scrambled or S1PR1 siRNA. Efficient S1PR1 downregulation (∼ 80±5% vs scramble) was achieved 24 h after transfection (Supplemental Figure 9). When cells were challenged with hypoxia, we found that hypoxia-induced barrier leakage was exacerbated in S1PR1 siRNA-treated cells (Figure 6A, 2.1±0.3 fold), compared to scrambled siRNA-treated (1.8±0.15 fold). To confirm the role of S1PR1 in barrier function using a pharmacological approach, we used the S1PR1 antagonist, W146 ^88^. Consistent with our data with genetic downregulation of S1PR1, acute pharmacological blockade of S1PR1 with W146 also heightened hypoxia-induced barrier leakage in HBMVEC compared to vehicle-treated cells (57% increase, Supplemental Figure 10). These data indicate that S1PR1 is an endogenous protective signaling pathway in the human cerebrovascular endothelium that limits hypoxia-induced barrier dysfunction.

**Figure 6.**
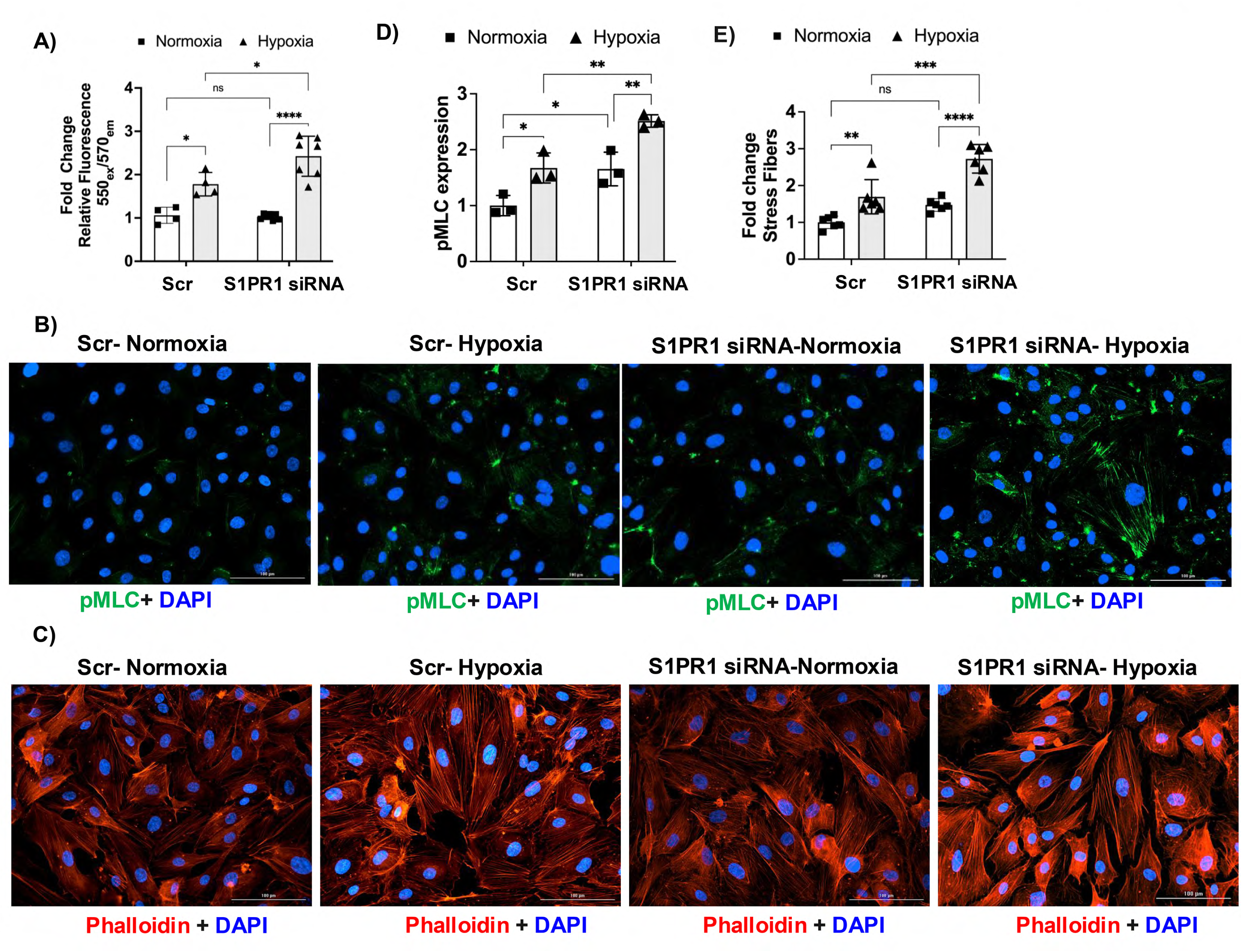
S1PR1 downregulation exacerbates hypoxia-induced primary human brain microvascular endothelial cell (HBMVEC) barrier dysfunction, phosphorylation of MLC and stress fiber formation. HBMVEC were transfected with scrambled (Scr) or S1PR1 siRNA and subjected to normoxia or hypoxia, as indicated in the methods section. A) Hypoxia-induced barrier leakage was exacerbated by S1pr1-knockdown compared to Scr transfected cells. Hypoxia-induced (B, D) phosphorylation of myosin light chain (pMLC) and (C, E) formation of stress fibers was heightened in S1pr1 siRNA transfected cells compared to Scr. B) IF staining for pMLC. C) Stress fiber formation was visualized by phalloidin staining. (B, C) Representative images from N=3-6 independent trials conducted in duplicates. Quantification was done using BioTek Lionheart automated imaging module Gen5. The data points and the mean ± SEM are plotted. *P < 0.05; **P < 0.01; ***P < 0.001; **** P < 0.0001; two-way ANOVA multiple comparisons followed by Tukey’s.

We also tested the role of S1PR1 in mouse cerebral endothelial barrier function *in vitro*. In the mouse endothelial brain cell line, bEnd3, downregulation of S1PR1 by siRNA or pharmacological blockade with the S1PR1 antagonist, W146, exacerbated hypoxia-induced leakage (3.7±0.8 fold and 1.9±0.23 fold, respectively, Supplemental Figure 11). Compared with HBMVEC, a more severe hypoxic challenge (6h) was required to induce robust barrier permeability (Supplemental figure 8B and D). These data indicate that in this *in vitro* system (hypoxia-induced barrier leakage), S1PR1 is an endogenous protective pathway that limits barrier dysfunction in the mouse brain endothelium.

Next, we aimed to determine the cellular and molecular mechanisms by which S1PR1 restricts hypoxia-induced barrier leakage in cerebral microvascular endothelial cells. We examined key cellular drivers of barrier dysfunction such as the phosphorylation of MLC, which mediates actomyosin assembly and cell contraction, the formation of stress fibers^82^ and caveolin-1 trafficking, a major driver of endothelial transcytosis^86^. Hypoxia increased the levels of p-MLC (1.67-fold, Figure 6B, D) along with the induction of stress fiber formation (1.69-fold, Figure 6C, E) and an increase in caveolin-1 levels (2.97-fold, Figure 7A, B). Downregulation of S1PR1 exacerbated hypoxia-induced phosphorylation of MLC (∼63%, Figure 6B, D) and stress fiber formation (∼73%, Figure 6C, E). Notably, downregulation of S1PR1 in normoxia significantly increased the expression of caveolin-1 (3.58-fold) compared to that in scramble siRNA-transfected HBMVEC (Figure 7A, B). The extent of induction of total Cav-1 by S1PR1 downregulation was similar to that induced by hypoxia (Figure 7B). Hypoxia-ischemia increases caveolin-1-dependent vesicular trafficking in the cerebral microvasculature, which mediates transcellular BBB permeability via the endosomal transcytosis pathway ^89^ ^86, 90^. We observed vesicle-like caveolin-1 positive structures in HBMVEC (Figure 7A). Further characterization of these Cav-1 positive vesicle-like punctates revealed that they were positive for the early endosomal markers Rab5a and EEA1, but not clathrin (Supplemental Figure 12), suggesting that they are endosomal vesicles ^86^. Next, we quantified the effect of hypoxia and S1PR1 expression on the number of endosomal Cav-1 and Rab5a positive vesicle-like punctates in HBMVEC. As shown in Figure 7C and D, S1pr1 siRNA knockdown induced an 80% increase in the number of Rab5a+ Cav-1+ vesicles during hypoxia. Taken together, these data indicate that S1PR1 signaling limits hypoxia-induced activation of MLC and stress fiber assembly, caveolin-1 endosomal transcytosis pathway, and endothelial barrier dysfunction in human brain microvascular endothelial cells.

**Figure 7.**
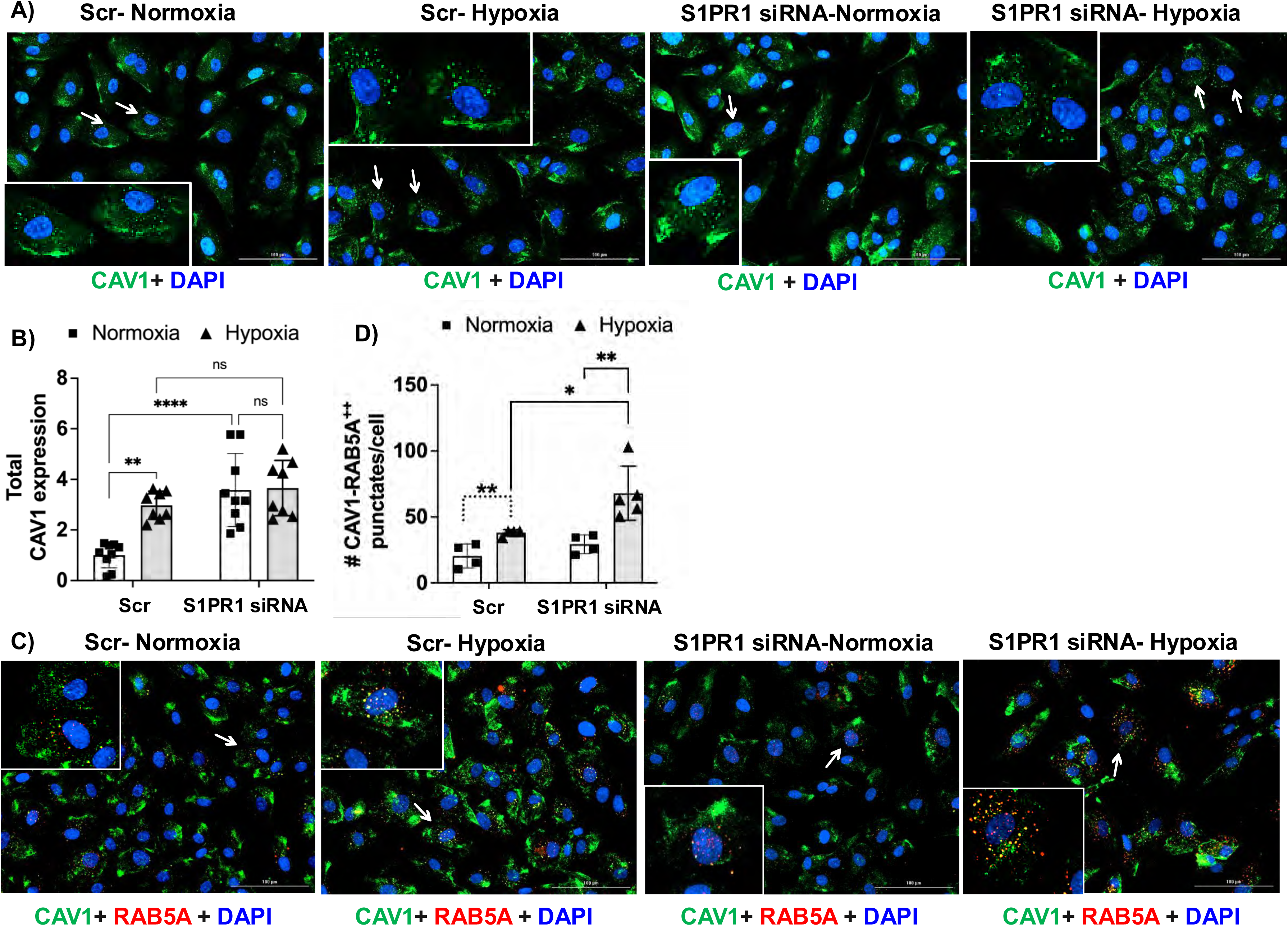
S1PR1 limits CAV1 expression and hypoxia-induced CAV1 localization with endosomal trafficking marker RAB5A in HBMVEC. HBMVEC were transfected with scrambled (Scr) or S1PR1 siRNA and subjected to normoxia or hypoxia. A) Immunofluorescence imaging of CAV1 in HBMVEC treated with S1pr1 or Scr siRNA subjected to normoxia or hypoxia. Inset: digital zoom (2x) of the cells marked with white arrows. B) Quantification of CAV1 immunofluorescence. Downregulation of S1PR1 with siRNA and hypoxia induces the levels of CAV1. Images were acquired at 20X magnification and analyzed using the BioTek Lionheart FX microscope and its image analysis module Gen5. C) Immunofluorescence imaging of CAV1 (green channel) and endosomal trafficking marker RAB5A (red channel) in HBMVEC. White arrows indicate examples of the CAV1 and RAB5A colocalization signal across the various experimental conditions as shown in the representative overlayed images. Inset: digital zoom (2x) of the cells marked with white arrows. D) Quantification of CAV1-RAB5A^++^ (double positive) signals. Hypoxia-induced CAV1 translocation to RAB5A endosomes was exacerbated in S1pr1 siRNA HBMVEC compared to Scr. The data points and the mean ± SEM are plotted. *P < 0.05; **P < 0.01; **** P < 0.0001; two-way ANOVA multiple comparisons followed by Tukey’s.

### Endothelial S1PR1 limits hypoxia-induced and ROCK-dependent MLC phosphorylation, stress fiber formation, Cav-1 endosomal distribution and barrier dysfunction

Given the protective role of S1PR1 in hypoxia-induced stress fiber formation and MLC phosphorylation, a downstream effector of Rho-ROCK, and the critical role of the cytoskeleton in vesicular trafficking, we aimed to determine the role of ROCK in these hypoxia-driven cellular processes. Pretreatment with the ROCK inhibitor Y-27632 (30 minutes) before the hypoxic challenge prevented MLC phosphorylation (Figure 8A, D), stress fiber formation (Figure 8B, E), and increased endosomal localization of cav-1 (Figure 8C, F) induced by S1PR1 knockdown under hypoxic conditions. Inhibition of these cellular events by blocking the Rho-ROCK pathway also prevented endothelial barrier dysfunction (Figure 8G). These data indicate that the blockade of S1PR1 signaling exacerbates hypoxia-induced barrier leakage, MLC phosphorylation, stress fiber formation, and cav-1 distribution to endosomes in a ROCK-dependent manner. Finally, we confirmed the role of S1PR1 in Rho-ROCK dependent hypoxia-induced endothelial barrier permeability by pharmacological inhibition of S1PR1, using the specific S1PR1 antagonist, W146 ^88^. Acute blockade of S1PR1 with W146 exacerbated hypoxia-induced barrier leakage in HBMVEC in a ROCK-dependent way (Supplemental Figure 13). Similarly, in bEnd3 cells the effects of pharmacological or genetic blockade of S1PR1 on hypoxia-induced leakage were counteracted by the ROCK inhibitor, Y-27632 (Supplemental Figure 14 A, B), confirming that S1PR1 restricts hypoxia-induced ROCK-dependent barrier dysfunction in mouse brain endothelial cells.

**Figure 8.**
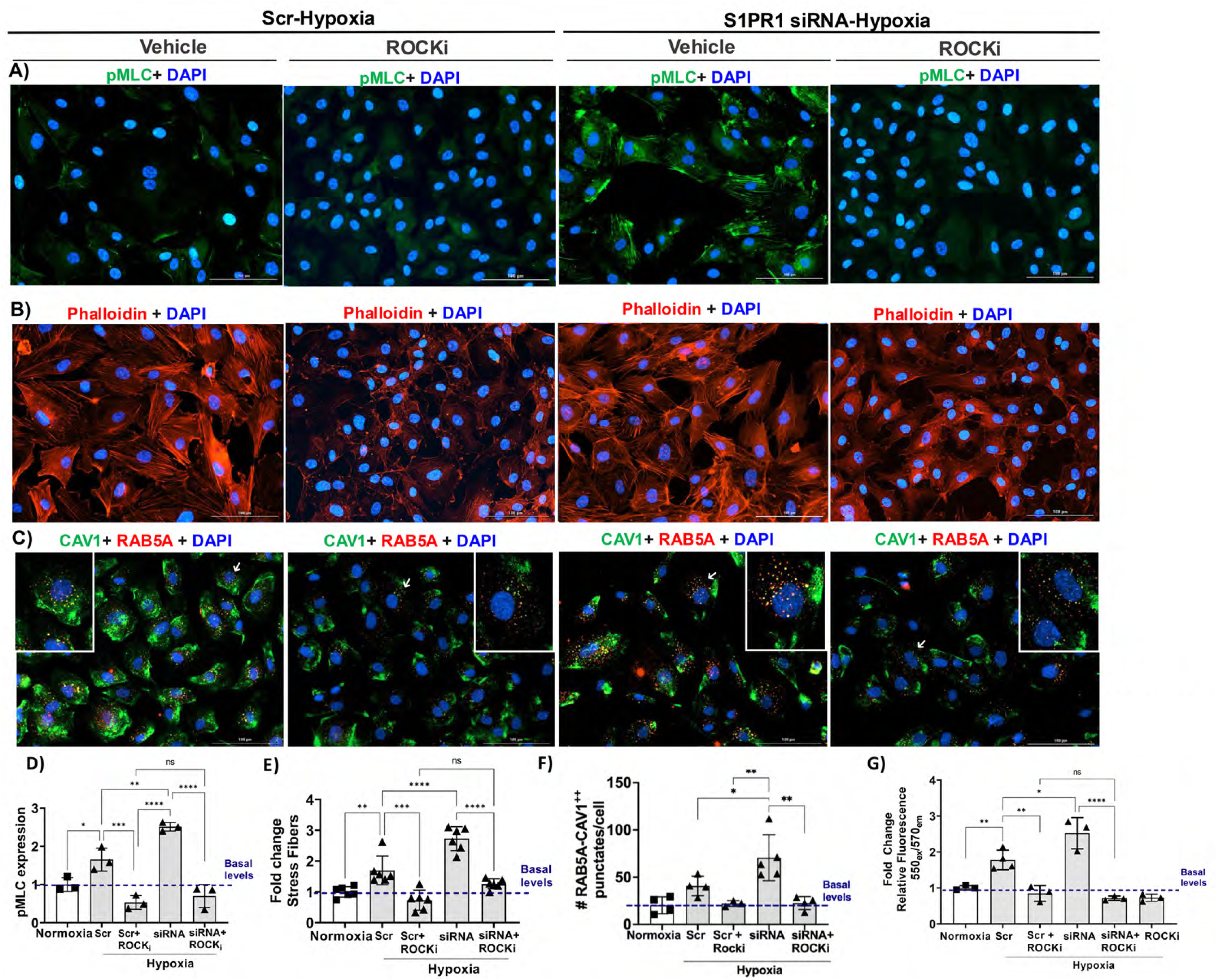
Hypoxia and S1PR1 knock down-induced MLC phosphorylation, stress fiber formation, CAV1 translocation to RAB5A+ endosomes and endothelial barrier dysfunction are ROCK dependent. HBMVEC were transfected with scrambled (Scr) or S1PR1 siRNA and subjected to hypoxia (30 min.) as indicated in the methods section. A, D) Hypoxia and S1pr1 downregulation induction of MLC phosphorylation is mitigated by treatment with the ROCK inhibitor Y-27632 (10µM) 30 minutes before hypoxic challenge. B, E) Hypoxia and S1pr1 siRNA-induced stress fiber formation was prevented by treatment with the ROCK inhibitor (Y-27632, 10µM). C, F) Inhibition of ROCK with 10µM Y-27632 prevents CAV1-RAB5A colocalization induced by S1pr1 downregulation in hypoxic HBMVEC. (G) Inhibition of ROCK reverses hypoxia and S1pr1 downregulation-induced barrier leakage in primary brain microvascular (HBMVEC) cells and restores endothelial barrier integrity comparable to basal (normoxia) state. Each datapoint represents an independent experiment performed in duplicated. Images were acquired at 20X magnification and analyzed using the BioTek Lionheart FX microscope and its image analysis module Gen5. The individual values and the mean ± SEM from N=3-6 trials are shown. *P < 0.05; **P < 0.01; ***P < 0.001, **** P < 0.0001 one-way ANOVA multiple comparisons followed by Tukey’s.

Altogether our *in vitro* studies indicate that in human primary brain microvascular endothelial cells, S1PR1 is a functional receptor which limits hypoxia-induced, ROCK-dependent cellular events that lead to barrier dysfunction, such as MLC phosphorylation, stress fiber formation and caveolin-1 assembly in endosomal vesicles. These in vitro data together with our data in the human brain showing the prevalence of S1PR1 in the cerebrovascular endothelium indicate that, like in mice, S1PR1 is a vasoprotective signaling pathway in the human cerebrovascular endothelium.

## SUPPLEMENTAL FIGURE LEGENDS

**Supplemental Figure 1. Glut-1 immunofluorescence in the S1PR1-eGFP knock in mouse brain.** Confocal analysis of S1PR1-eGFP fluorescence and Glut-1 immunodetection in mouse cortex. Notice the expression of S1PR1-eGFP (green channel) throughout the brain parenchyma but also in the cerebrovascular endothelium (labeled by immunofluorescence for Glut-1, endothelial marker, red channel). A) Representative pictures are shown. N=5-6. Sections were captured using an FluoView FV10i confocal microscope (Olympus, Japan) (original magnification, x 60). Notice the expression of S1PR1-eGFP in microvessels (white arrows). B) High power view (digital zoom) of selected area in panel A. Scale bar 20μm.

**Supplemental Figure 2. MAP-2 immunofluorescence in brain sections in the S1PR1-eGFP knock in mice.** S1PR1-eGFP fluorescence confocal analysis in grey matter (A, cortex and B, hippocampus) and white matter areas (C, D, internal capsule) of the mouse brain. Notice that S1PR1-eGFP signal is localized in the dendrites and neuronal processes (MAP-2 positive). Scale bar 20μm. D) High power view (digital zoom) of inset shown in panel C. Image shows localization of S1PR1-eGFP around the microtubules. Scale bar 10μm. Sections were imaged by using an FluoView FV10i confocal microscope (Olympus, Japan), (original magnification, x 60). Representative pictures are shown. N=5-6

**Supplemental Figure 3. Expression of S1PR1-eGFP and Nissl stain in brain sections in the S1PR1-eGFP knock in mouse.** S1PR1-eGFP fluorescence confocal analysis in grey matter (A, cortex and B, hippocampus) and white matter areas (C, corpus callosum) of the S1PR1-eGFP knock in mouse brain. Neuronal somas are stained with NeuroTrace® 530/615 Red Fluorescent Nissl (red channel). Note that S1PR1-eGFP is expressed in neurons and localized around the soma, mainly in the neuropil (A, B) and neuronal processes (C). Sections were imaged by using a FluoView FV10i confocal microscope (Olympus, Japan), (original magnification, x 60). Scale bar 20μm. Representative pictures are shown. N=5-6.

**Supplemental Figure 4. Expression of S1PR1-eGFP and GFAP immunofluorescence in brain sections in the S1PR1-eGFP knock in mice.** S1PR1-eGFP fluorescence confocal analysis in white matter areas (A, B, corpus callosum, C, internal capsule) of the mouse brain. Arrows indicate fibrous astrocytes and processes positive for S1PR1-eGFP. B) Digital zoomed image of two fibrous astrocytes positive for S1PR1-eGFP (arrows). C) Digital zoomed image of an astrocyte negative for S1PR1-eGFP (arrow). Sections were imaged by using an FluoView FV10i confocal microscope and a 60x objective (Olympus, Japan) Scale bar 20μm (A), 10μm (B,C). Representative pictures are shown. N=5-6.

**Supplemental Figure 5. S1PR1 mRNA is the most abundant S1PR transcript in brain and highly enriched in cerebral microvessels. A)** S1PR mRNA levels in the mouse brain. Quantitative reverse transcription and polymerase chain reaction (RT-qPCR) demonstrates that S1PR1 transcript is predominant over the other S1PR. **B)** Relative S1PR1 mRNA levels in isolated cerebral microvessels compared to whole brain shows that S1PR1 is highly enriched in cerebral microvessels. The individual values and the mean ± SEM are shown. N=3-6. *p<0.05.

**Supplemental Figure 6. SAH induces S1PR1 protein levels in cerebral microvessels.** Western blot analysis for S1PR1 in brain microvessels 24 after subarachnoid hemorrhage (SAH). Microvessels were isolated and S1PR1 levels were detected by western blot (*n* = 4). A) Immunoblot representative image. B) Western blot quantification was conducted using Image J. Fold change of S1PR1 protein levels in sham *versus* SAH (normalized by β-actin). Individual values and mean ±SEM are shown. \*\**P* < 0.01, t test.

**Supplemental Figure 7. Endothelial S1PR1 limits SAH-induced BBB leakage in females and middle-aged (12-month-old) mice.** A) Albumin BBB leakage, assessed by Evans Blue Dye extravasation, 24 hours after sham or SAH surgery in *S1pr1^flox/flox^*and *S1pr1^iECKO^* young female mice and B) *S1pr1^flox/flox^*and *S1pr1^iECKO^* male middle-aged mice. Evans blue dye was circulating for 3 hours. *Y* axis shows relative fluorescence units (R.F.U.) per gram of tissue and normalized by sham. The individual values and mean ± SEM are shown. *n* = 8 - 19. \**P* < 0.05, \*\**P* < 0.01, \*\*\**P* < 0.001, \*\*\*\**P* < 0.0001, one-way ANOVA followed by Tukey’s test. Discontinuous line indicates t test.

**Supplemental Figure 8. Hypoxia induces human and mouse brain endothelial barrier dysfunction and upregulation of caveolin-1.** A, B) Endothelial barrier leakage (4.4KDa MR-Dextran) in primary human brain microvascular endothelial cells, HBMVEC (A), and mouse cortical brain endothelial cells, bEnd.3 (B), subjected to the indicated times of hypoxia or normoxia. A, B) The mean ± SEM of n=3-6 experiments are shown. C) HBMVEC barrier permeability after 30-minute exposure to hypoxia. D) bEnd.3 barrier permeability after 6 hour-exposure to hypoxia. E, F) Induction of hypoxia marker, vascular endothelial growth factor-a, *VEGF* mRNA and caveolar protein, caveolin-1 (Cav1) mRNA in HBMVEC after 30-minute exposure to hypoxia. G, H) Induction of *Vegf* mRNA and caveolin-1 mRNA in bEnd.3 cells after 6 hour-exposure to hypoxia. *P < 0.05; **** P < 0.0001; one-way ANOVA multiple comparisons followed by Tukey’s correction. (C-H) Individual values and the mean ± SEM are shown. Each data point represents an independent experiment. *P < 0.05; **P < 0.01; ***P < 0.001; **** P < 0.0001; Unpaired t-test.

**Supplemental Figure 9. siRNA knockdown of S1pr1 in primary human brain microvascular endothelial cells.** Depletion in the levels of S1pr1 transcripts in HBMVEC. Cells were treated with scrambled (Scr) or S1pr1 siRNA for 24h and exposed to hypoxia. qPCR of total RNA collected from scrambled control and siRNA treated cells reveals a significant depletion (80%) in S1pr1 mRNA levels. The individual values and the mean ± SEM normalized by Scr are shown. N = 3 independent experiments. **P < 0.01, Unpaired t-test.

**Supplemental Figure 10. Pharmacological inhibition of S1PR1 induces brain endothelial barrier permeability.** Treatment of primary human brain microvascular endothelial cells, HBMVEC, with the S1PR1 specific antagonist W-146, significantly induces endothelial barrier leakage. Cells grown in transwell filters, treated for 30 min. with 10µM of W-146 compound and challenged with hypoxia for additional 30min. Next, barrier dysfunction was determined by quantifying 4.4 kDa TRITC-Dextran leakage from the top to the bottom chamber. Data is presented from n=3 independent trials conducted in duplicates. Values presented as mean ± SEM normalized by vehicle hypoxia with **P < 0.01, Unpaired t-test.

**Supplemental Figure 11. Blockade of S1PR1 by siRNA knockdown or pharmacological inhibition exacerbates endothelial barrier permeability in mouse brain endothelial cells.** A) Knockdown of S1pr1 using siRNA in mouse brain endothelial cells (bEnd.3). bEnd.3 were grown until confluence in normal culturing conditions and transfected with scrambled or mixed pool of S1pr1 siRNA. 24 hours later they were subjected to hypoxia for 6 hours. Thereafter, cells were harvested, total RNA was extracted as described in the methods section and processed for quantitative real-time PCR. Data were collected from N=6 independent trials. B) Knockdown of S1rp1 induces endothelial barrier dysfunction. bEnd.3 cells were grown in transwell filter plates and transfected with S1pr1 or scrambled siRNA. Barrier permeability was determined by 4.4KDa TRITC-conjugated dextran leakage from the top to the bottom compartment after 24 h of hypoxic challenge. Data were collected from n=3 independent trials conducted in duplicates. C) The S1PR1 antagonist W146 increases barrier leakage under hypoxic conditions. bEnd.3 cells were pre-treated with 10µM W-146 for 30 minutes before hypoxic challenge. 24 h later, barrier leakage was determined. Individual values presented as mean ± SEM normalized by scrambled or vehicle hypoxia. *P < 0.05; **** P < 0.0001, Unpaired t-test.

**Supplemental Figure 12. Cav-1 positive vesicle-like punctates co-localize with endosomal markers Rab5a and EEA1 in HBMEC.** After 30 min. of hypoxic challenge, HBMVEC cells were fixed and processed for IF analysis for Cav-1, and the vesicular trafficking markers A) Rab5a, B) early endosome marker EEA1 and C) clathrin. CAV1+ vesicle-like structures were also positive for RAB5A (A) and EEA1 (B) (two examples are labeled with white arrows). In contrast, clathrin staining exhibited a distinct pattern of localization that of CAV1, as it can be seen in the overlay image. All images were acquired at 20X magnification and analyzed using the BioTek Lionheart FX microscope and its image analysis module Gen5. Inset: digital zoom (2x) of the cells marked with white arrows.

**Supplemental Figure 13. Pharmacological inhibition of Rho kinase abrogates S1PR1 blockade-induced cerebral endothelial barrier permeability during hypoxia.** The S1PR1 antagonist W-146 (10µM) induces barrier dysfunction as seen by induced small molecule leakage in HBMVEC (∼1.5-fold). Treatment with the ROCK inhibitor Y-27632 (ROCKi, 10μM) restores endothelial barrier integrity as seen by marked reduction in dextran leakage. The individual values and the mean ± SEM normalized by vehicle hypoxia are plotted. *P < 0.05; **P < 0.01; **** P < 0.0001 one-way ANOVA multiple comparisons followed by Tukey’s.

**Supplemental Figure 14. Pharmacological inhibition of ROCK abrogates cerebral endothelial barrier permeability induced by S1PR1 blockade.** A) 24 h after transfection with control (Scr) or S1pr1 siRNA, bEnd.3 cells were treated with the ROCK inhibitor (ROCKi, Y-27632, 10μM) for 30 min. and subjected to hypoxia for 6 hours. Genetic knockdown of S1pr1 induces small molecule leakage (∼4-fold vs Scr control), which was abrogated by Y-27632 treatment. B) Pharmacological inhibition of S1pr1 by W-146 (10µM) induces barrier dysfunction (∼2-fold vs vehicle, Veh), which was abrogated by ROCK inhibition with 10μM Y-27632. Each datapoint represents an independent experiment. The individual values and the mean ± SEM, normalized by vehicle scrambled hypoxia are plotted. *P < 0.05; **P < 0.01; **** P < 0.0001 one-way ANOVA multiple comparisons followed by Tukey’s.

