## Supplemental Figures for "Endothelial Sphingosine-1-Phosphate Receptor (S1PR) 1, a Functional S1PR in the Human Cerebrovascular Endothelium, Limits Blood Brain Barrier Permeability and Neuronal Injury following Subarachnoid Hemorrhage in Mice"

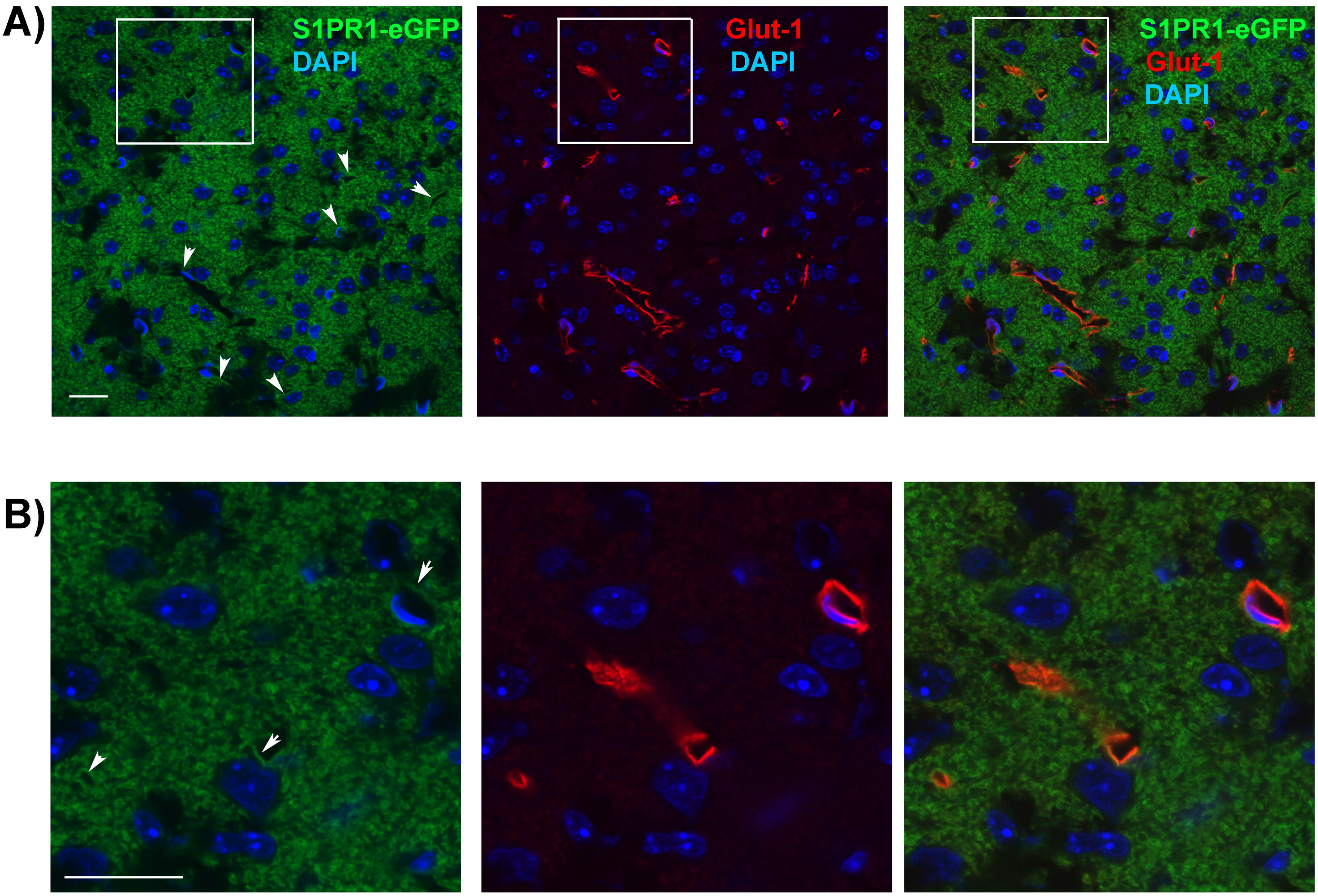


### Supplemental Figure 1

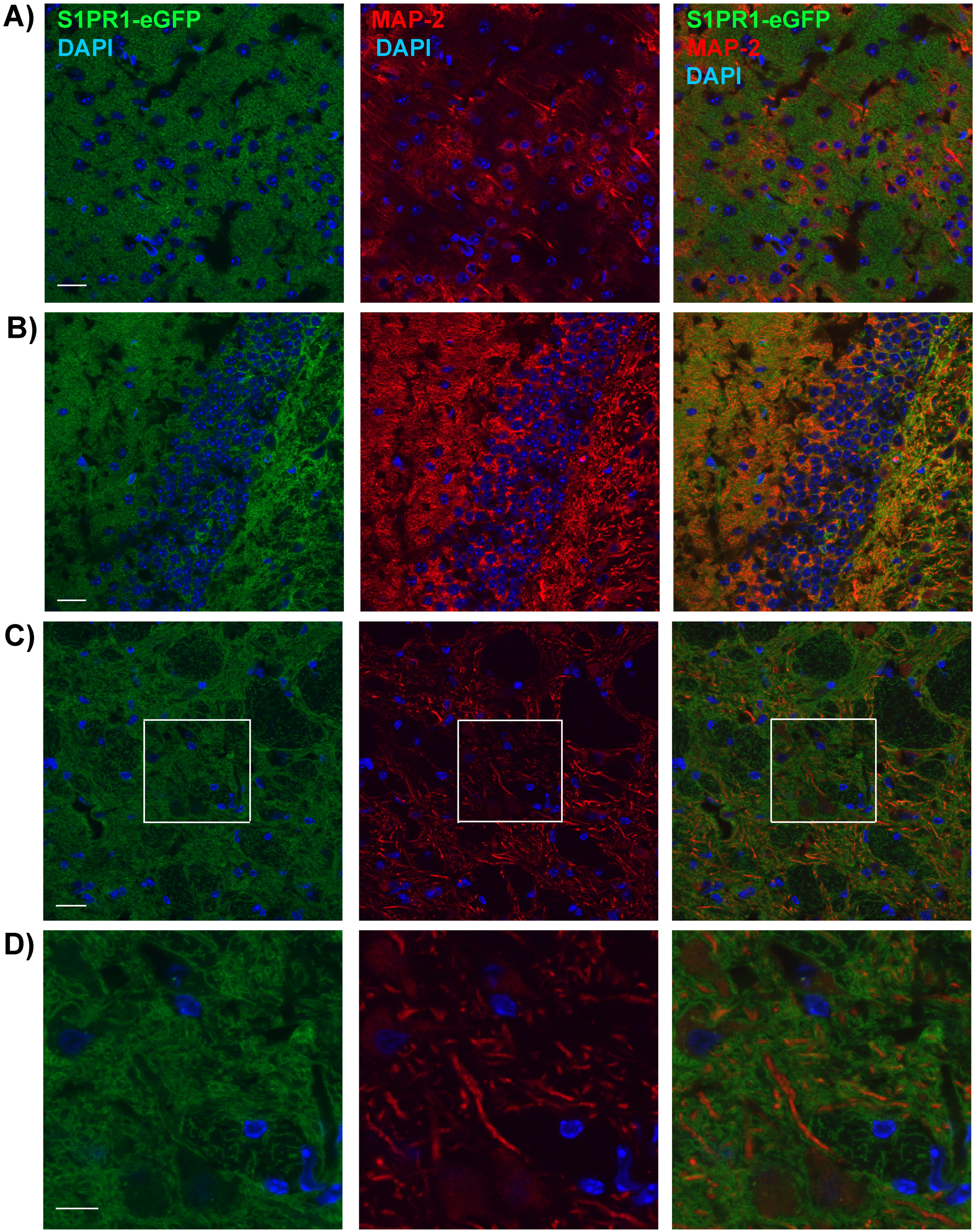


**Supplemental Figure 2**

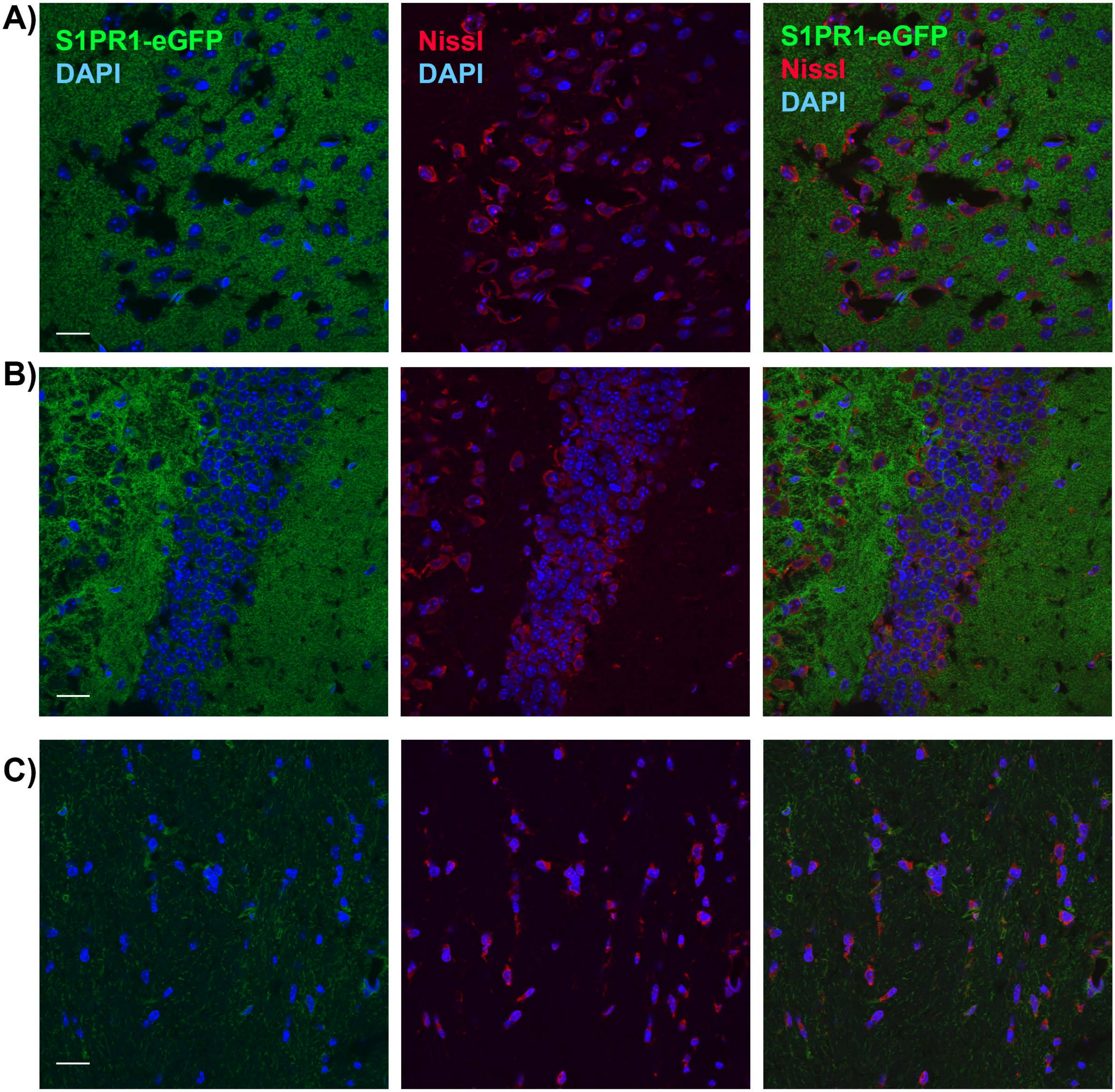


**Supplemental Figure 3**

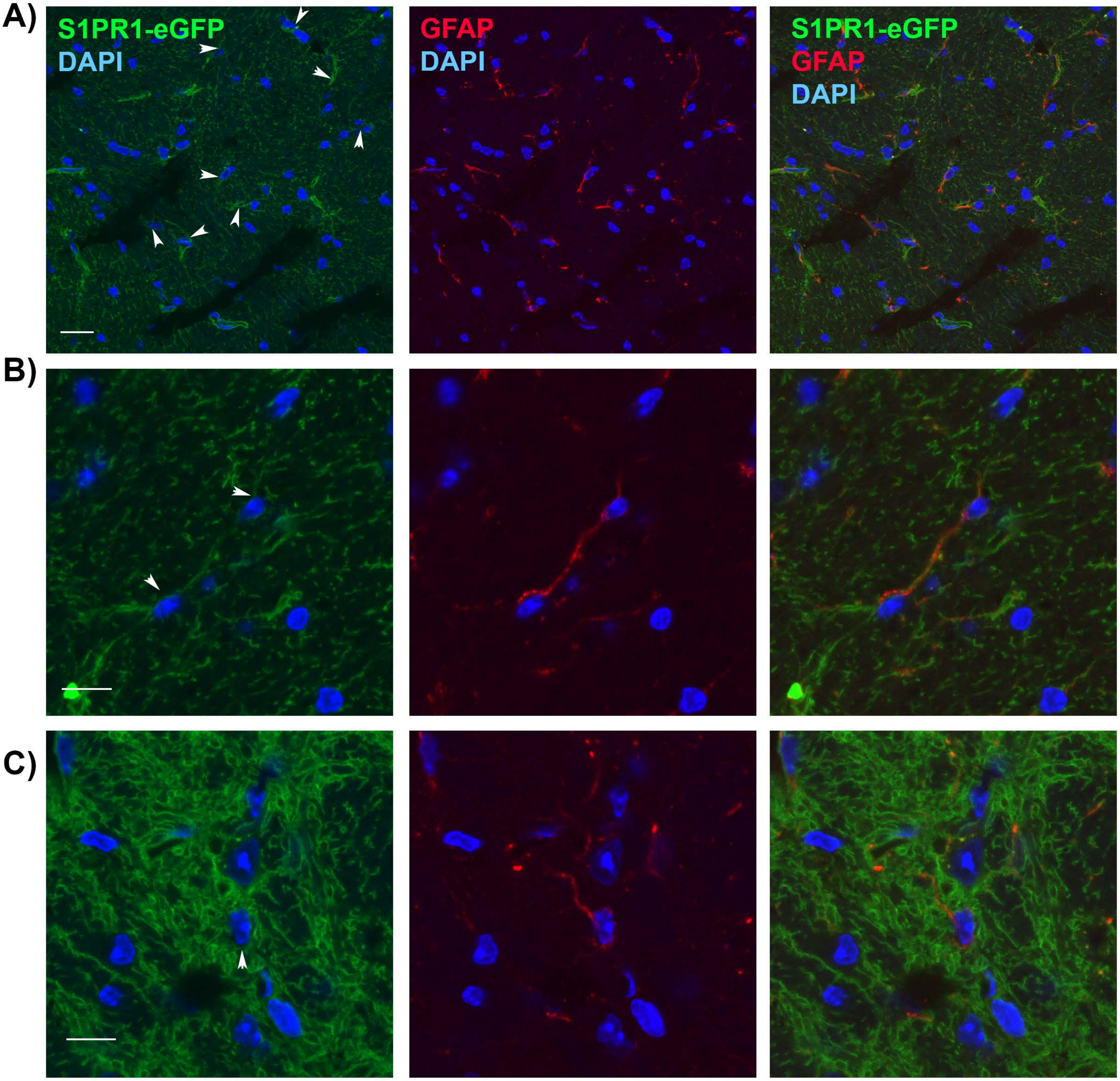


**Supplemental Figure 4**

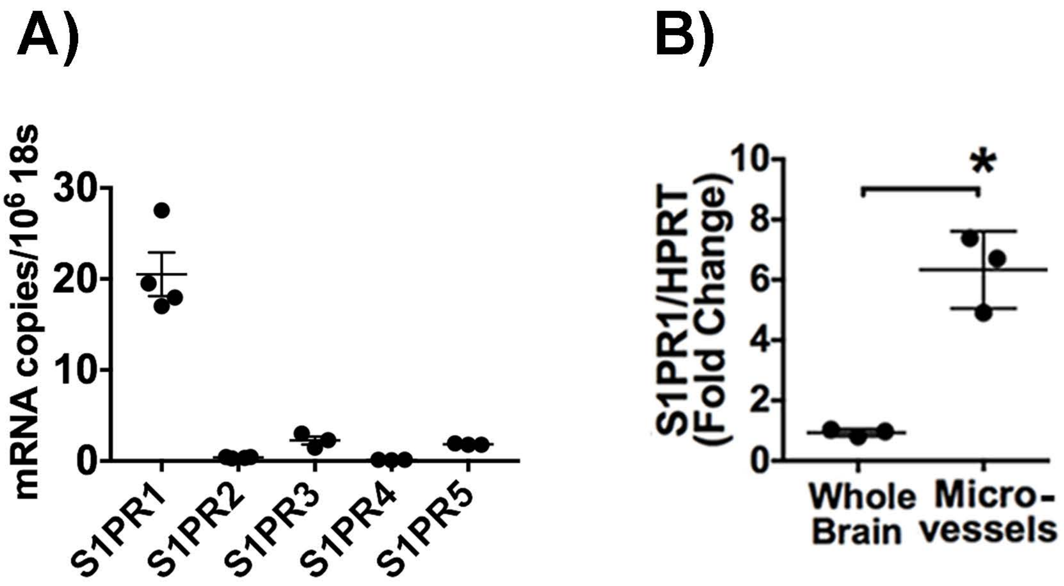


### Supplemental Figure 5

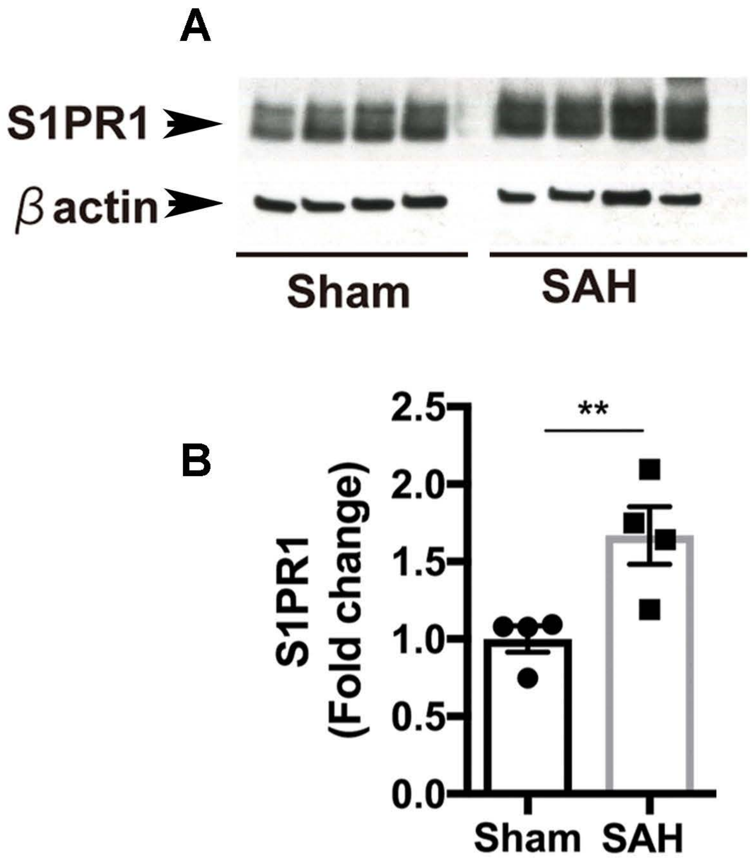


Supplemental Figure 6

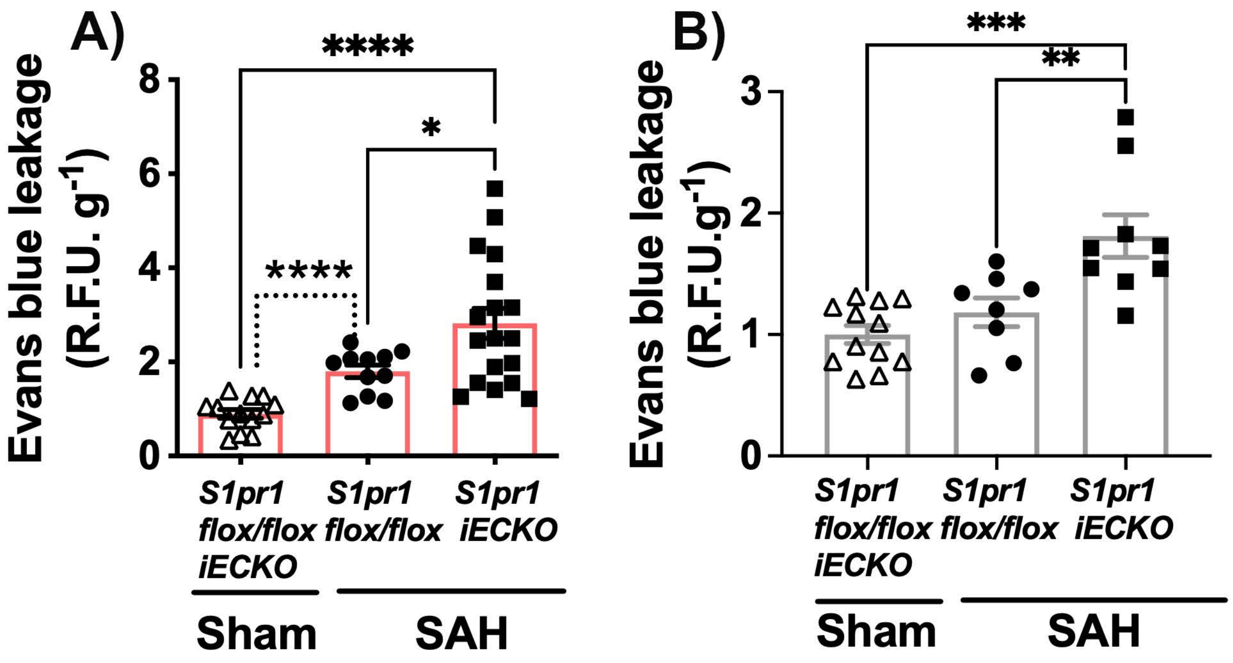


### Supplemental Figure 7

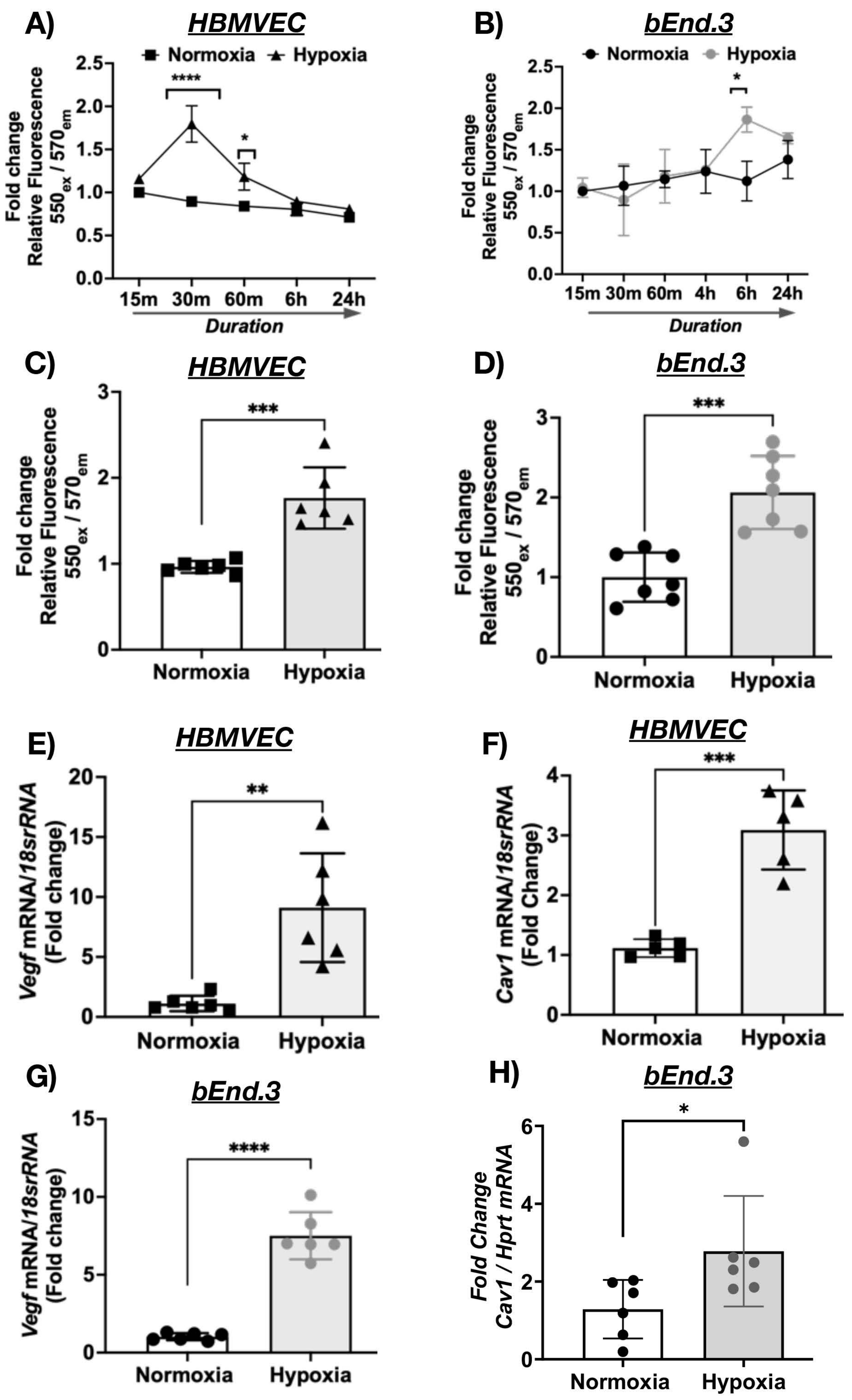


Supplemental Figure 8

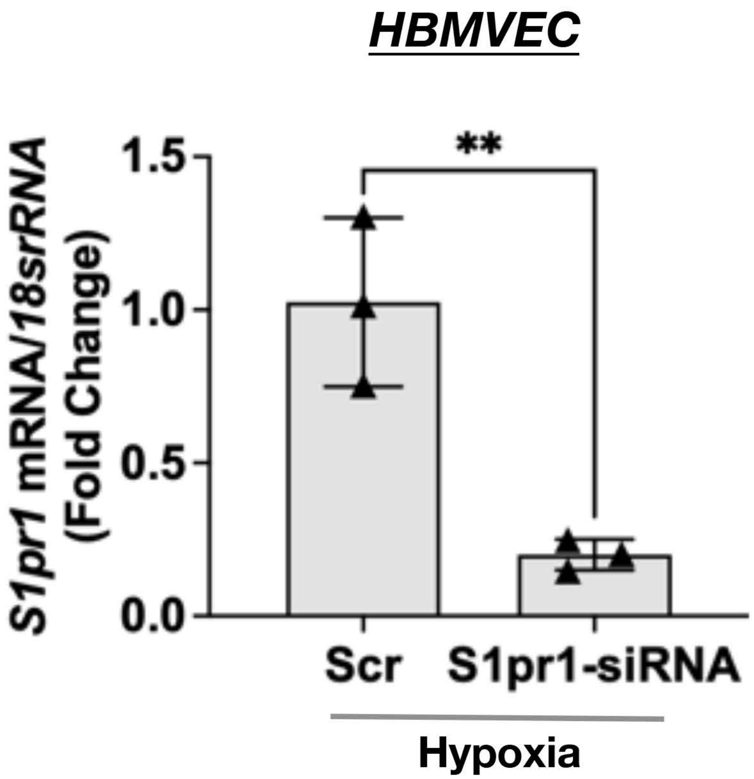


Supplemental Figure 9

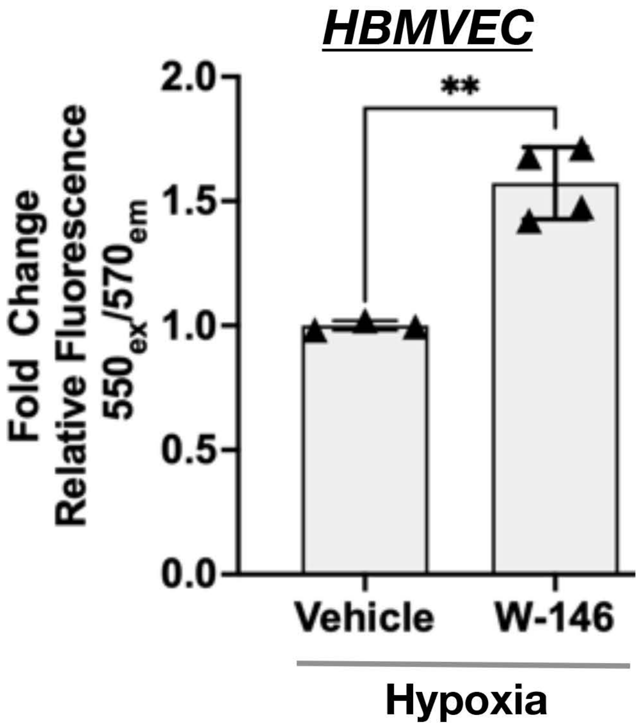


Supplemental Figure 10

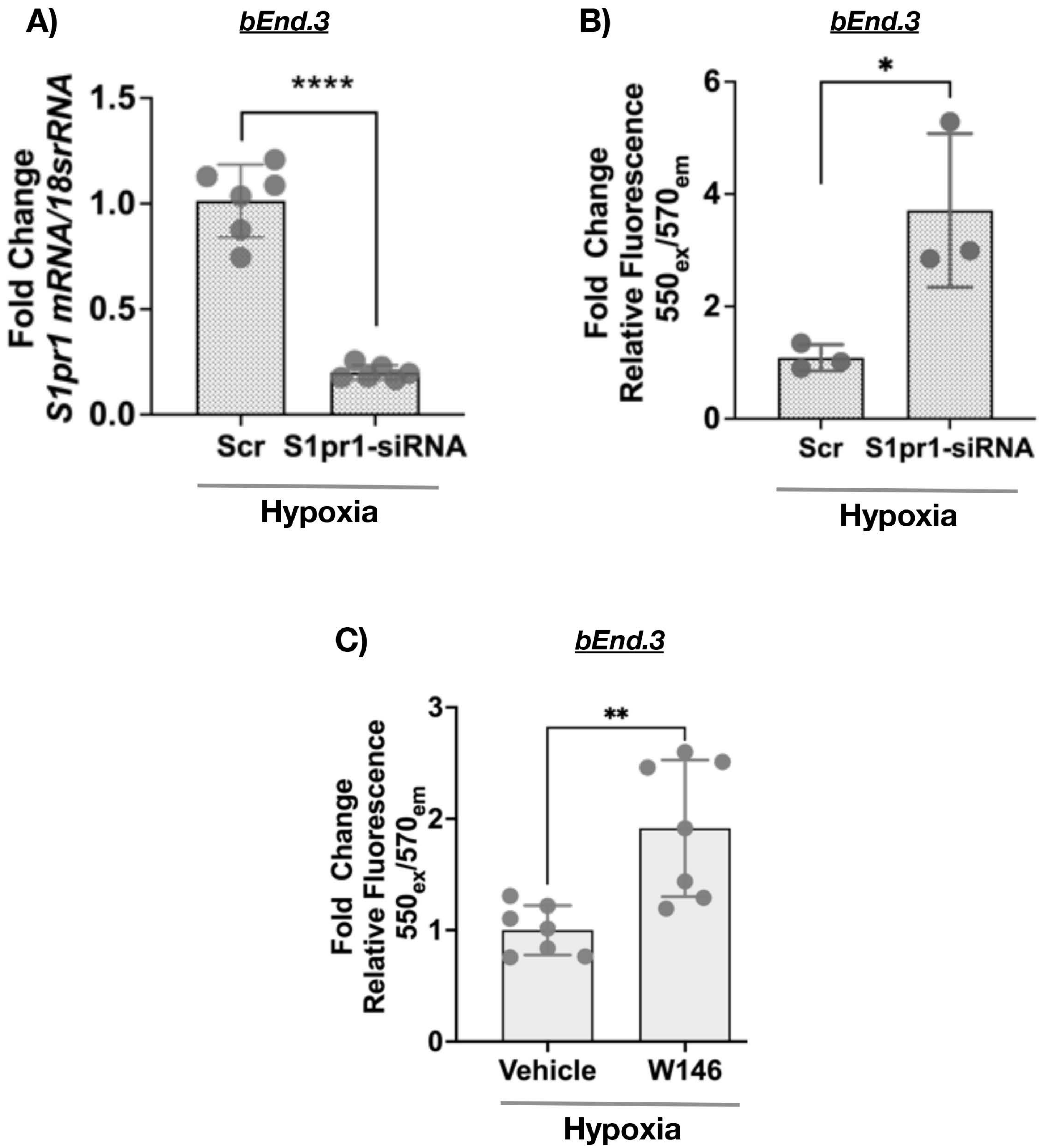


**Supplemental Figure 11**

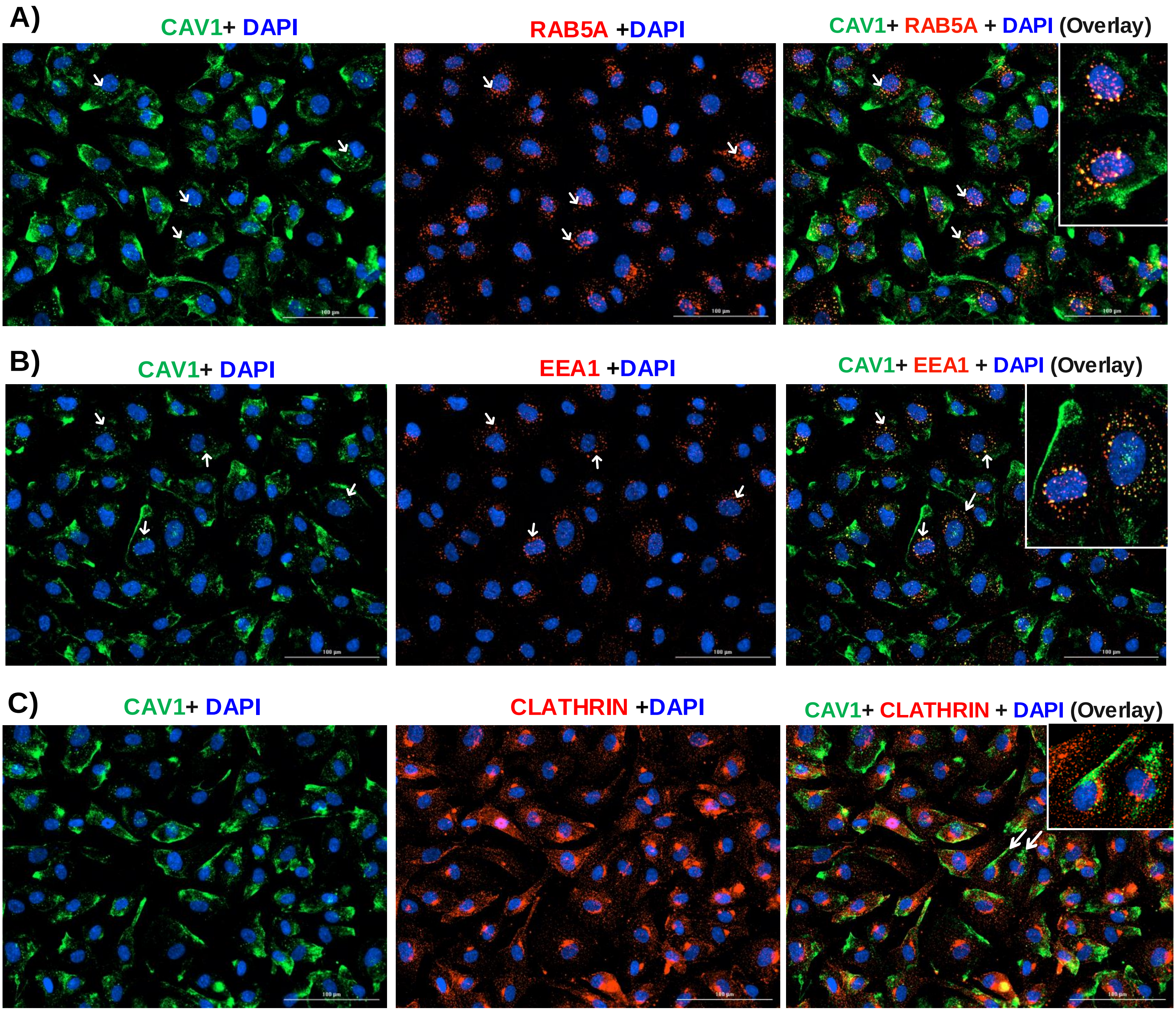


**Supplemental Figure 12**

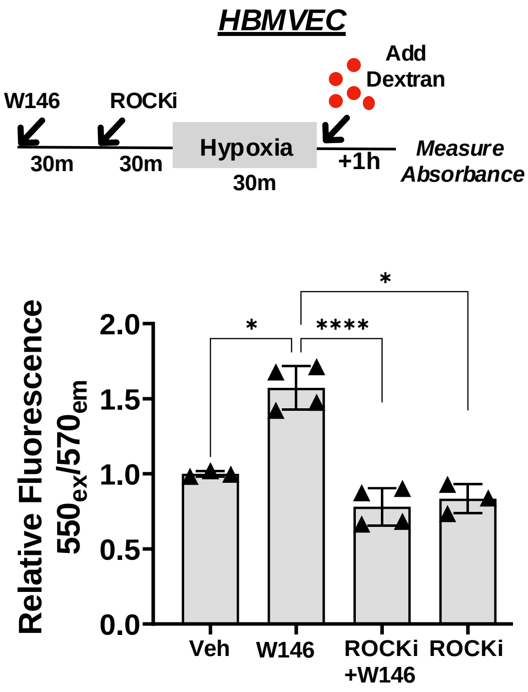


### Supplemental Figure 13

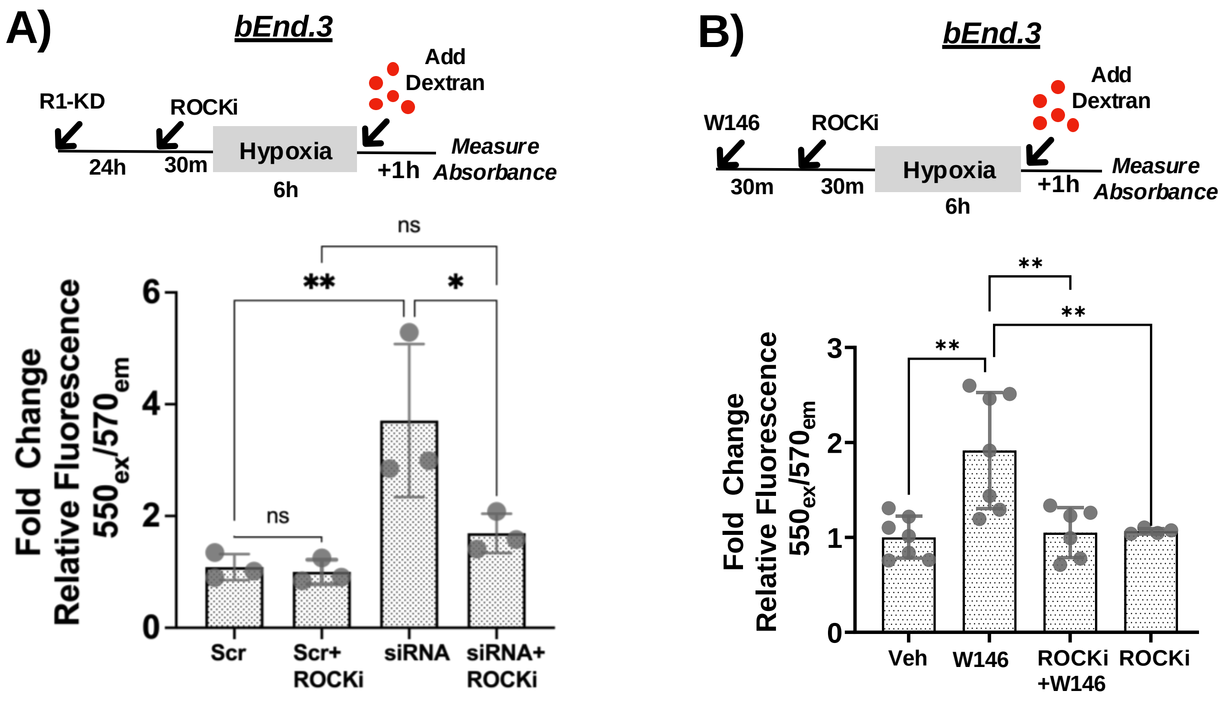


### Supplemental Figure 14
